# Basic-Leucine-Zipper Transcription Factors Regulate Selective Molecular Phenotypes in Regulatory T Cells During IL-2-Induced Activation

**DOI:** 10.1101/2025.02.25.638325

**Authors:** Joyce Tse, Xuan Liu, Ittai Eres, Tracy Yamawaki, Matt Kanke, Marisela Killian, Junedh Amrute, Ashutosh Chaudhry, Anupama Sahoo, Wenjun Liu, Cheng-Yuan Kao, Xin Luo, Jiamiao Lu, Daniel Lu, Songli Wang, Scott Martin, Chi-Ming Li

**Affiliations:** Functional Genomics of Research Technologies, Amgen Global Research, South San Francisco, California, United States of America; Center for Research Acceleration by Digital Innovation (CRADI), Amgen Global Research, South San Francisco, California, United States of America; Center for Cardiovascular Research & Division of Cardiology, Department of Medicine, Washington University School of Medicine, Saint Louis, Missouri, United States of America; Inflammation Research, Amgen Global Research, South San Francisco, California, United States of America; Amgen R&D Postdoctoral Fellows Program, South San Francisco, California, United States of America

**Author notes:** Equal contribution (Joyce Tse, Xuan Liu).

**Keywords:** regulatory T cells, Treg activation, Treg heterogeneity, basic leucine zipper, multi-omics, chromatin accessibility, transcription factor motif, chromatin remodeling, epigenomics, transcriptomics

## Abstract

Regulatory T (Treg) cells have long been recognized as modulators of immunological tolerance and homeostasis. Previously, we used scRNA-seq to reveal significant Treg heterogeneity in response to IL-2-induced activation. Herein, we leveraged enrichment analyses, as well as bulk and single-nucleus multi-omics in splenic and lung Tregs, to uncover and confirm the importance of transcription factors (TFs) and chromatin remodeling in Treg activation. Multiple bZIP TF motifs showed increased chromatin accessibility post IL-2 treatment, with correlated transcriptional changes resembling Th1 and Th2 molecular phenotypes, further confirmed by spatial ATAC-seq. By combining gene perturbation and CUT&RUN assays before and after Treg stimulation, we show that bZIP TFs, such as BATF and BACH1, are critical to IL-2-induced Treg activation, coordinating epigenetic and transcriptional changes that selectively drive T-helper phenotypes and metabolic pathways.

## Introduction

Since the 1970s, T cells have been known to be capable of suppressing experimentally-induced autoimmune diseases in rodents (Gershon and Kondo, 1970). Further investigations led to the discovery of CD25 (interleukin-2 receptor α chain, IL-2Rα) as a highly expressed molecular marker of suppressive CD4^+^ T cells, which are now known as regulatory T (Treg) cells (Sakaguchi et al., 1995). Tregs are cells of the adaptive immune system that act as suppressors of conventional T cells and are essential for immunological tolerance and homeostasis. In 2003, Forkhead box protein 3 (FOXP3) was identified to be a master regulator of Treg development, specification and function (Fontenot et al., 2003, Hori et al., 2003). Despite the significance of FOXP3, it is insufficient by itself to specify the complete Treg transcriptional program. Given evidence of FOXP3-independent signatures that regulate Treg function (Hill et al., 2007), it is worth interrogating the regulation of both FOXP3 positive and negative CD4^+^CD25^+^ cells by applying different activation factors for Treg cells, such as IL-2.

IL-2 is crucial for Treg cell development, differentiation, survival, proliferation and suppressive function. It also acts on pro-inflammatory lymphocytes and plays a central role in the amplification of immune responses. It has been shown that IL-2-induced Treg activation is associated with increased chromatin accessibility, as assessed by bulk <u>A</u>ssay for <u>T</u>ransposase <u>A</u>ccessible <u>C</u>hromatin with sequencing (ATAC-seq) (Moro et al., 2022). It identified TFs (*Stat5*, *Jund*) and chromatin remodelers (*Batf*, *Ezh2*) involved in Treg activation. Others have also found *PRDM1/Blimp-1*, *NF-κB*, *Batf*, and *Ezh2* as important transcriptional regulators of Treg activation or maintenance of activation (Ogawa et al., 2018, Oh et al., 2017, Itahashi et al., 2022, DuPage et al., 2015). Interestingly, *PRDM1/Blimp-1* knockout in gut Treg cells leads to IL-17 expression with loss of suppressor function to promote intestinal inflammation, indicating potential T-helper 17 cell phenotype of such Treg cells (Ogawa et al., 2018). In fact, Treg cells can, under certain transcription factor programs, differentiate into “specialized” Treg cells with T-helper 1, 2, 17 or T-follicular helper (Th1, Th2, Th17, or Tfh) phenotypes (Trujillo-Ochoa et al., 2023). However, the link between chromatin remodeling and transcription programing for Treg associated T-helper cell characteristics is not well studied.

Interestingly, the outcome of IL-2 signaling is dependent upon the composition of differential affinity IL-2 receptor (IL-2R) chains and IL-2 expression levels (Hernandez et al., 2022). Treg cells express a high affinity trimeric IL-2R comprising of CD25 (IL-2Rα), CD122 (IL-2Rβ) and CD132 (IL-2Rγ). This allows preferential Treg IL-2R binding and signaling at low IL-2 dosage. Recently, IL-2 engineering has led to enhancement of target specificity and prolonged activity. To better understand non-transient Treg-specific functions, recombinant forms of IL-2, including IL2-IgG2a Fc fusion protein (Fc-IL-2) or further half-life extended form of Treg-specific mutant IL-2, which was generated as a IgG2a Fc fusion protein with the additional amino acid mutations to promote binding to CD25 (IL-2 mutein; IL-2M), have been well-characterized and currently used in the field. The half-life of IL-2M is on the order of days, compared to hours for recombinant IL-2. Our previous single-cell RNA sequencing (RNA-seq) analysis identified various Treg cell states, including resting, primed, and proliferative states as well as a subset of activated Treg cells that co-express *Il1rl1* and *Tnfrsf9,* encoding surface proteins, ST2 and 4-1BB, respectively, and demonstrate high immuno-suppressive function after Fc-IL-2 stimulation (Lu et al., 2020), suggesting that ST2/4-1BB expression may provide an efficient way to separate cell states while investigating Treg activation. Interestingly, a substantial number of TFs and chromatin DNA binding factors that modulate transcription and/or chromatin remodeling are present as the gene signatures of these highly activated states (Lu et al., 2020).

To characterize the transcriptomic and epigenomic landscape of Treg cell activation, we analyzed sorted ST2^+^/4-1BB^+^ and ST2^-^/4-1BB^-^ Treg cells by using bulk RNA-seq and ATAC-seq. Single-nucleus ATAC-seq analysis revealed dynamic alterations to chromatin accessibility primarily by bZIP TF (basic leucine zipper transcription factor) motifs along the activation spectrum. Bulk and single-nucleus transcriptomes corroborated comparable activity of bZIP TF-mediated Treg activation upon IL-2 treatment. To further interrogate these factors, we performed CUT&RUN experiments to assess the binding regions of bZIP TFs of interest, ultimately identifying the target genes of bZIP TF regulation. We also employed gene perturbation studies to validate the importance of specific bZIP TFs in Treg cell activation and chromatin accessibility. Finally, we performed extensive expression profiling and spatial ATAC-seq analysis on Treg marker genes, revealing molecular phenotypes of T-helper-like subsets. In summary, we provide systematic evidence that the superfamily of bZIP TFs, along with increases in chromatin accessibilities of their motifs are critical regulators that regulate multiple pathways during Treg cell activation.

## Results

### ST2^+^/4-1BB^+^ Tregs demonstrate transcriptional signatures of activated Treg cells

To benchmark the highly activated Treg cells expressing ST2 and 4-1BB over the subset that negatively express both of these two surface proteins (herein referred to as “resting” Tregs), we leveraged such surface co-expression to define and isolate the highly activated spleen Treg subpopulations by fluorescence-activated cell-sorting (FACS) after CD4^+^CD25^+^ magnetic enrichment. As seen in the workflow in Figure 1A and S1, after four days of murine IL-2M (mIL-2M) treatment, activated and resting Treg cells were subjected to bulk RNA-seq and, later, ATAC-seq analysis. Gene expression profiling analysis identified distinct transcriptional expression patterns. The 3775 upregulated genes and 3492 downregulated genes (adjusted P-value <0.05, |Log2(FC)|>0.25 in order to capture more subtle effects) while comparing the activated and resting Treg cells are listed in Tables S1 & S2. Signature markers of activation (*Il10*, *Clta4*, *Ccr8*, and *Tnfrsf* family genes) and of resting states (*Satb1*, *Tcf7*, *Sell, Ccr7*) were identified in the list of up- and down- regulated differentially expressed genes (DEGs), respectively (Figures 1 B and C). Gene Set Enrichment Analysis (GSEA) highlighted IL-2/STAT5 signaling (Figure 1D) and showed enrichment for components in the mTORC1 pathway, MYC targets, and various regulators involved in cell proliferation, including E2F targets, G2M checkpoint factors, and mitotic spindle genes (Figure S2). In addition, Ingenuity Pathway Analysis (IPA) identified enrichment of cell cycle genes and antigen processing or presentation genes expressed by activated Treg cells, confirming the transcriptional signature of mIL-2M-induced proliferation and activation in ST2^+^/4-1BB^+^ Tregs (Figure 1E). GO analysis based on Molecular Function revealed enrichment of chromatin-DNA binding genes in ST2^+^/4-1BB^+^ Treg cells (Figure 1F) in agreement with our previous finding. We further aligned the bulk RNA-seq upregulated DEGs to the single-cell-derived cluster DEGs of the two ST2^+^/4-1BB^+^ states and found that more than 80% of *Ass1^+^* and 60% of *Itgb7^+^* Treg cluster genes from our reference scRNA-seq were detected in the current bulk assay. However, the cluster DEGs only accounted for ∼13% of the bulk transcriptome, suggesting that ST2^+^/4-1BB^+^ Tregs represent a diverse and rich source of data worth interrogating with further multi-omics analysis to understand Treg activation (Figure S3).

**Figure 1.**
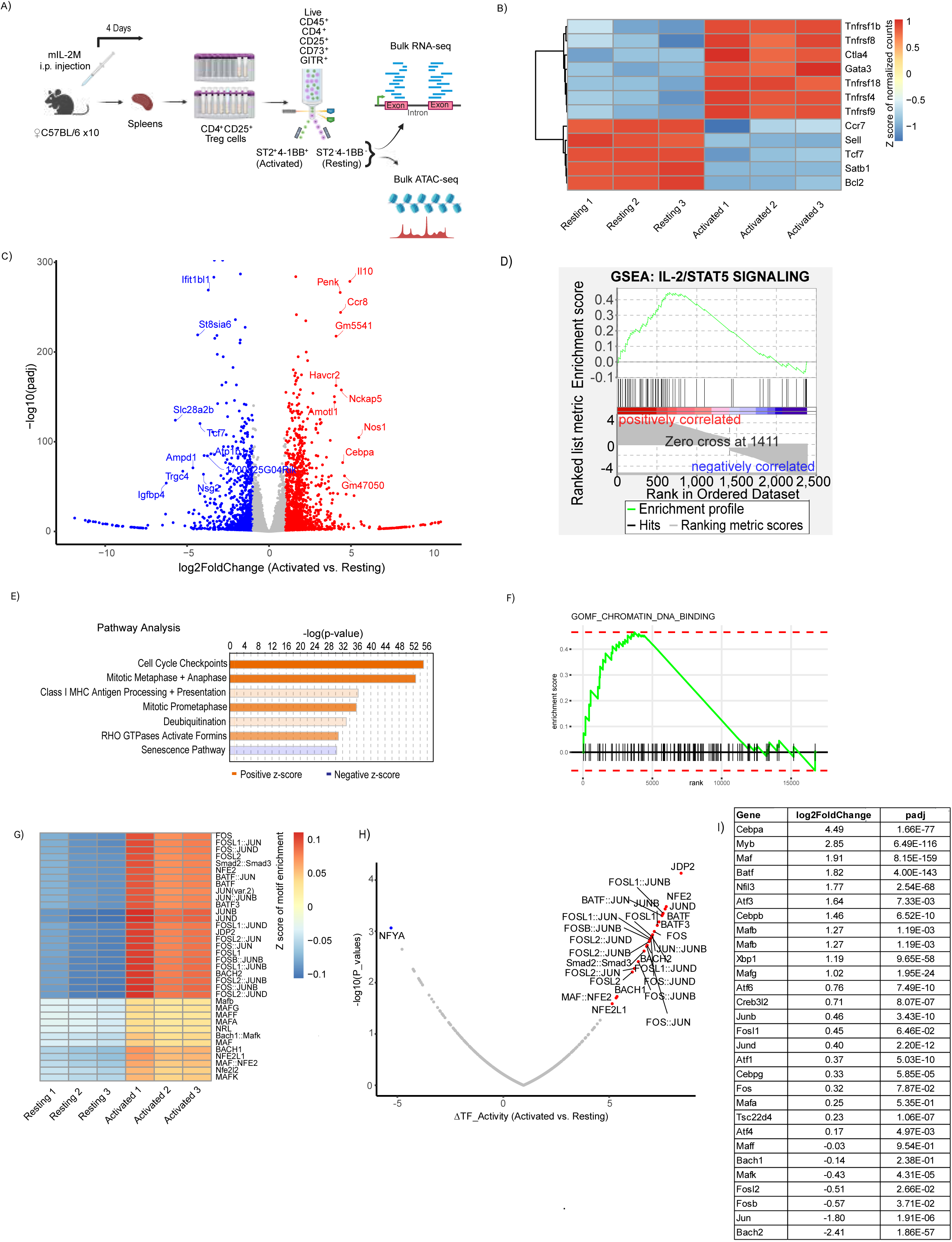
Benchmarking bulk multi-omics of resting and highly activated Treg cells identifies distinctive epigenetic profiles. (A) Experimental workflow for mouse splenic Treg cell isolation for bulk RNA-seq and bulk ATAC-seq. (B) Heatmap of normalized gene counts for specific resting and activation state marker genes for each sample. (C) Volcano plot of up- (red) and down-regulated (blue) differentially expressed genes with cutoffs of log2FoldChange > 1 or log2FoldChange < −1, respectively, and adjusted p-value < 0.05. Top 10 most up- and down-regulated genes with adjusted p-value < 10^-5^ were labeled. (D) Enrichment of IL-2/STAT5 signaling in activated Treg cells by Gene Set Enrichment Analysis. (E) Pathway analysis of all significant genes (adjusted p-value < 0.05). (F) Enriched chromatin-DNA binding in activated Treg cells by gene ontology analysis of all significantly (adjusted p-value < 0.05) upregulated differentially expressed genes. (G) Heatmap of enriched DNA binding motifs. The deviation (75% quantile) and significant difference between resting and activated states (adjusted p-value < 0.002). (H) A table showing the differentially expressed basic leucine zipper (bZIP) genes (activated versus resting Treg cells), along with the expression fold change and the corresponding adjusted p-value.

### IL-2 induced activated Treg cells are enriched in bZIP transcription factor motifs

To investigate the epigenomic landscape associated with Treg cell states, the chromatin accessibility profiles were assessed by ATAC-seq, resulting in the identification of distinct patterns of chromatin accessibility peaks between ST2^-^/4-1BB^-^ and ST2^+^/4-1BB^+^ subpopulations. Among the peaks with increased accessibility in the ST2^+^/4-1BB^+^ cells, we observed *Batf*, a pioneer TF, chromatin remodeler and critical regulator in Treg cell activation. ATAC peaks near or within the *Batf* gene appeared more accessible, with higher amplitude and greater total number of peaks, in ST2^+^/4-1BB^+^ Treg cells (Figure S4A). Additionally, the downstream targets of BATF, like *Il10* (whose expression was significantly upregulated in activated Treg cells) and *Il1rl1* (encoding the ST2 protein), were both more accessible in the ST2^+^/4-1BB^+^ Treg cells (Figures S4B and S4C) (Delacher et al., 2020b). We annotated differentially accessible regions with their closest genes (Table S3) and also observed that certain marker genes of the resting state became less accessible in the ST2^+^/4-1BB^+^ Treg cells compared to the ST2^-^/4-1BB^-^ Treg cells. In particular, the cell migration marker *Ccr7* (Figure S4D) and resting markers, *Satb1* (Figure S4E) and *Sell* (Figure S4F), exemplified this effect. The correlation of chromatin accessibility with RNA expression for these renowned Treg markers provides another layer of confidence motivating further exploration of the chromatin peaks between the ST2^-^/4-1BB^-^ and ST2^+^/4-1BB^+^ Treg cells, representing the resting and activated states, respectively.

To assess the interplay between differential accessibility and potential TF activity, we performed motif analysis on the ATAC peaks to identify DNA binding sequences recognized by specific types of TFs and predominantly enriched in either resting or activated states (Table S4). Strikingly, in addition to the SMAD2/3 complex motif in response to TGFβ signaling, the enriched motifs largely represented consensus binding sequences of bZIP TFs, which could be categorized into at least four classes (Figure 1G). The first class of the enriched motifs belongs to the FOS family (FOS, FOSB, FOSL1, FOSL2) or the JUN family (JUN, JUNB, JUND) in activated cells. FOS and JUN members heterodimerize to form AP-1 complexes, which are known to be involved in T cell activation (Jain et al., 1992). Next, the motifs of ATF family members, including BATF transcriptional activators, were also enriched in the activated state. The third class represents the enriched motifs of small MAF TFs (MAFF, MAFG, MAFK), which lack a transactivation domain. These TFs need to form homo- or hetero-dimers within the small MAF family, or with other types of transactivation factors (Tsuchiya et al., 2015) or with large MAF TFs (MAFA, MAFB and NRL, which contain a transactivation domain), in order to show enhanced associated transactivation functions. Lastly, the motifs of TFs belonging to the cap “n” collar (CNC) family, including NFE2 and BACH1/2, were also enriched.

To confirm the association of TF-DNA binding activity with motif accessibility, TF footprint analysis was performed on the ATAC peaks. We first examined the footprints of a number of bZIP TF motifs, including BATF, BATF3, BACH1, BACH2, JUNB, JUND, FOS, FOSL1, and FOSL2, which indicated increased binding activity in activated Treg cells (Figure S5). In order to systematically identify active TF binding sites and changes in TF binding activities between cell states, we compared the resting and activated Tregs to obtain TFs with significant changes using HINT-ATAC (Li et al., 2019). As expected, a number of bZIP TFs were significantly upregulated in the activated cells (Figure 1H). We further classified these TFs as activators or repressors according to their mode of action using the diffTF algorithm (Berest et al., 2019). Positively correlated bZIP TFs, in which increased chromatin accessibility corresponded to increased gene expression levels, were denoted as “activators,” including BATF, FOSL1, JUNB and JUND (Figures S6A to S6D). Negatively correlated bZIP TFs, in which increased chromatin accessibility corresponded with decreased gene expression levels, were denoted as “repressors,” which included BACH1/2 and FOSL2 (Figures S6E and S6F). Judging from the bulk DEG list of activated Treg transcriptomes, most bZIP transcriptional activators showed significant upregulation of expression, but several identified repressors were downregulated (Figure 1I).

### Single-nucleus multiome analysis confirms bZIP family gene regulation and signatures in activated Treg states compared to other cell states

In order to decipher chromatin accessibility and transcriptomic profiles at single cell resolution for various Treg cell states, the 10x Genomics single-nucleus multiome assay was performed on CD4^+^CD25^+^ Treg cells purified from mouse spleen or lung tissues after 4-days of mIL-2M stimulation (Figure S7A). Based on inferred gene activity and label transfer of the reference Treg scRNA-seq data (Lu *et al*. 2020), Treg cell activation states were annotated in both snATAC-seq and snRNA-seq datasets (Figure S7B). The resulting annotated chromatin accessibility profiles were subsequently used for motif and TF footprint analysis. We integrated our snATAC-seq and snRNA-seq data by comparing chromatin accessibility versus expression level of various candidate genes.

The heterogeneity of Tregs determined by previous scRNA-seq was recapitulated by both annotated snATAC-seq and snRNA-seq (Figures 2A and 2B). We observed global changes of accessible chromatin peaks between treated groups (Figure 2C), resulting in a reproducible pattern of differential chromatin DNA footprints that could be more or less accessible in either control or mIL- 2M-treated samples as shown in Table S5. We also observed apparent transitions of differential chromatin accessibility profiles between different Treg cell states (Figure 2D). Table S6 lists the accessible or inaccessible genomic regions shown in the cell state heatmap. Treg cells in a primed state (C2/*Ms4a4b^+^*), had a similar chromatin accessibility profile to resting (C1/*Satb1^+^*) Treg cells, with additional chromatin accessibilities that overlapped with the proliferative (C6/*Stmn1^+^*) state. A portion of accessible peaks in the proliferative state overlapped with the two activated states, which were previously annotated as *Ass1^+^* and *Itgb7^+^*(a.k.a. C4 and C8, respectively) Treg cells. Both activated states have further accessible peaks that overlapped partially with the peaks in memory state (C3/*Nr4a2^+^*) Treg cells. Interestingly, memory state Treg cells also shared chromatin peaks with the resting and primed states (Figure 2D). Overall, our data suggests that the majority of accessible chromatin loci in the resting and primed states become closed in the activated states, whereas we observed increased chromatin accessibility at regions originally packed or closed in resting Treg cells. Taken together, these transitions suggest that epigenomic trajectory may change over Treg activation.

**Figure 2.**
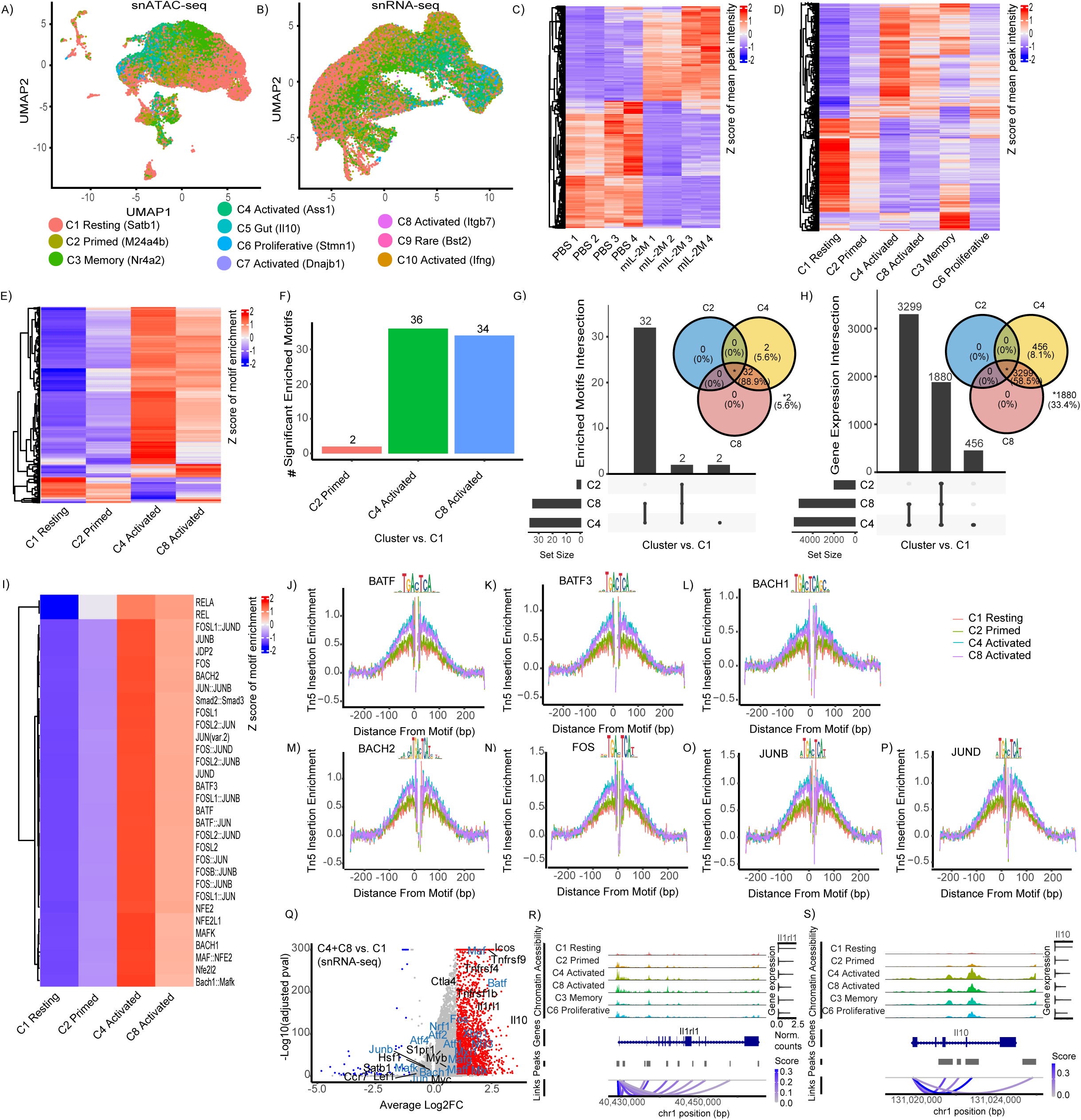
Highly activated Treg cells are enriched in basic leucine zipper transcription factor motifs and DNA binding activities by single-nuclei multi-omics. (A) UMAP plot generated from snATAC-seq and colored by predicted cell states. (B) UMAP plot generated from snRNA-seq and colored by predicted cell states. (C) Heatmap of chromatin accessibility for detected genomic regions for each pseudobulk sample. Read counts for peaks were aggregated by sample. The quantile of sum of peak counts were calculated, and the peaks whose sum of counts between the 20th and 90th percentile were kept for PBS and mIL-2M pseudobulk samples separately. The peak counts were normalized by DESeq2 median of ratios method. The top 5000 most variable peaks were selected and visualized by z-scores in a heatmap. (D) Heatmap of chromatin accessibility for detected genomic regions for various cell activation states using pseudobulk samples. Read counts for peaks were aggregated by sample. The quantile of sum of peak counts were calculated, and peaks falling between 20th and 90th percentile of all pseudobulk samples were kept. The peak counts were normalized by DESeq2 median of ratios method. The top 5000 most variable peaks were selected and visualized by Z-scores in heatmap. (E) Heatmap of enriched motifs for C1 resting, C2 primed and C4/C8 activated states, with cutoffs adjusted p-value < 0.05 and average difference > 0.1. The average motif activity score in each cell state was calculated and the z-score normalization was applied across cell states for visualization in the heatmap. (F) The number of enriched motifs of C2 primed and C4/C8 activated states relative to C1 resting state, with cutoffs adjusted p-value < 0.05 and average difference > 1. (G) The overlap of enriched motifs between cell states. (H) The overlap of genes downstream to enriched motifs. Promoter regions were defined as 2 kbp upstream to transcription start site (TSS) of coding genes. Peaks containing motifs and overlapping with promoter regions were selected. The closest genes whose coding sequence (CDS) downstream to the selected peaks were pulled out as downstream genes. (I) Heatmap of enriched motifs for cell states, with more stringent cutoffs adjusted p-value < 0.05 and average difference > 1. (J-P) The footprinting profiles of selective basic leucine zipper transcription factors depicting the difference between C4 and C8 activated states to the C1 resting and C2 primed states. (Q) Volcano plot showing the differentially expressed genes between activated (C4 + C8) and resting (C1) cell states with bZIP genes labeled in blue. Genome browser view showing the peak intensities for genes *Il1rl1* (R) and *Il10* (S) in the C4 and C8 activated states compared to all other Treg cell states.

We performed motif analysis on our snATAC-seq peaks, revealing that activated states were enriched with a greater number of motifs compared to resting and primed states (Figure 2E and Table S7). As shown in Figure 2F, the C4/*Ass1^+^* and C8/*Itgb7^+^* activated states had 36 and 34 significantly enriched motifs (relative to C1/*Satb1^+^*), respectively, whereas the C2/*Ms4a4b^+^*primed state only had 2 significantly enriched motifs (Table S8). Interestingly, these 2 enriched motifs, REL (MA0101.1) and RELA (MA0107.1), were also found in both of *Ass1^+^* and *Itgb7^+^* activated states (Figure 2G), suggesting these motifs could have a role in stability or maintenance of Treg cell activation. Indeed, the number of downstream genes regulated by these 2 motifs represent a drastically large network of 1880 genes out of 5635 cluster-defining DEGs (33.4%) (Figure 2H). Notably, out of a total of 36 unique enriched motifs, 34 (94.4%) were shared between *Ass1^+^* and *Itgb7^+^* activated states, suggesting shared roles in maintaining the characteristics of Treg cell activation. As shown in Tables S9 and S10, respectively, these motifs impact 5179 downstream genes (91.9%). There were also 2 enriched motifs unique to the C4/*Ass1^+^* activated state, NFκB (MA0105.4) and MAFG (MA0659.2), suggesting these may be important in initiating the C4/*Ass1^+^* activation state or transitioning from the resting or primed state in mIL-2M treated Tregs (with 456 downstream genes; 8.1%).

Assessing statistical significance of enriched motifs in the *Ass1^+^*and *Itgb7^+^* activated states revealed a list that comprised primarily of four classes of bZIP TF motifs at cutoff thresholds of adjusted p-value < 0.05 and average difference > 1 (Figures 2I and Table S8), in concordance with the findings from bulk ATAC-seq analysis. It is worth noting that REL and RELA motifs were also enriched in both *Ass1^+^*and *Itgb7^+^* activated states although they were not detected in the ST2^+^/4-1BB^+^ activated Treg cells in bulk assays. This could likely be a result of REL and RELA motifs also enriched in *Ms4a4b^+^*primed state that are part of the ST2^-^/4-1BB^-^ Tregs. Interestingly, by decreasing the cutoff threshold (from average difference > 0.5 to > 0.25 without changing adjusted p-value < 0.05), Kruppel-Like Factor (KLF), TBX and RUNX TF family motifs started to appear in the activated states as well (Figure S8), suggesting that transcriptional regulation involved by bZIP TFs may be more active than that of the other TF classes during IL-2-induced Treg activation. In order to further assess cell state-specific motif activity, the footprints of multiple bZIP TFs were individually analyzed across cell states. Similar to the bulk assessment above, we found that the motifs of BATF, BATF3, BACH1, BACH2, FOS, JUNB and JUND in the activated states from the single-nuclei assay showed increased binding activity relative to resting and primed states (Figures 2J to 2P).

We annotated Treg activation cell states in the snRNA-seq data and identified DEGs in the activated Treg cells compared to the resting ones, with a volcano plot indicating the significantly (adjusted p-value < 0.01) and highly upregulated (average log2 fold change ≥ 1, red dots) and downregulated (average log2 fold change ≤ −1, blue dots) DEGs (Figure 2Q). Reassuringly, in activated Treg cells, we observed upregulation of bZIP TF superfamily genes (e.g. *Maf*, *Batf*, *Fos*, *Nrf1*, *Xbp1*, *Atf1*, *Atf2*, *Atf4*, *Mxi1*, *Myb*, *Myc*, *Maff*, *Mafg*, *Mafk*, *Mlx*, *Bach1*, *Jun*, *Junb*, *Hsf1*, blue text) and activation genes (e.g. *Icos*, *Tnfrsf9*, *Tnfrsf4*, *Tnfrsf1b*, *Il1rl1*, *Il10*, *Ctla4*, *Nfil3*) as well as downregulation of resting genes (e.g. *Satb1*, *Lef1*, *Ccr7*). We additionally generated volcano plots based on the “gene activity” assay derived from snATAC-seq, labeling bZIP TF genes with resting and activated marker genes, or the top 15 DEGs for both this and the RNA assay from snRNA-seq (Figure S9). Interestingly, as one of the top upregulated genes of cell activation, *Il1rl1* shows stronger chromatin accessibility in activated states, in its gene body as well as in upstream and downstream regions (Figure 2R). Similarly, *Il10* displays increased chromatin accessibility and gene expression in the activated states from single-nuclei multi-omics, corroborating findings in activated Treg cells from bulk and multi-omics assays (Figure 2S). Taken together, our multi-omics data provide evidence that, due to dynamic changes in chromatin accessibility in activated Treg cells, various bZIP TFs may be driving the distinct expression patterns of transcriptional signature genes between Treg cell states, and hence appear critical for mIL-2M-induced Treg activation.

### Lung Treg cells demonstrate a similar bZIP TF motif landscape to that identified in splenic counterparts post mIL-2M stimulation

We extended our investigation to Treg cells purified from non-lymphoid lung tissues by using the single-nuclei multi-omics assays. Similarly to our annotation of splenic single-nuclei multi-omics data, we visualized the landscapes of lung Treg snATAC-seq or snRNA-seq, respectively based on gene activities and gene expression levels, as UMAP plots in Figures 3A and 3B. The identities of most Treg cell states matched between spleen and lung although the frequencies of their *Nr4a2*^+^ activated state differed (Figure S10). Overall, lung Treg cell comparisons between PBS control and mIL-2M treated groups yielded distinct profiles of chromatin accessibility but comparable to the ones generated by using splenic snATAC-seq analysis (Figure 3C and Table S11).

**Figure 3.**
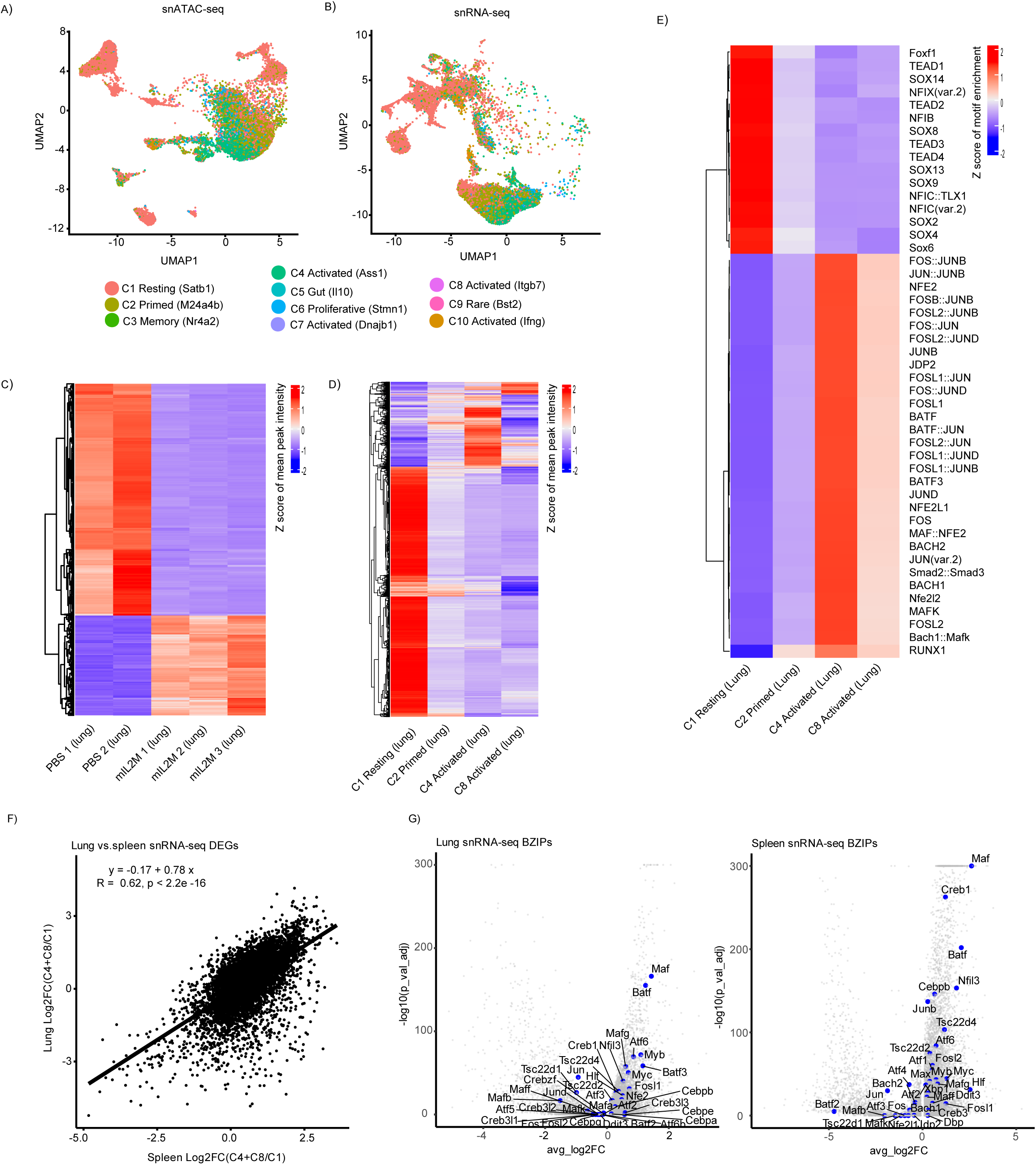
Non-lymphoid tissue lung is also enriched in basic leucine zipper motifs. (A,B) UMAP plots for multi-state lung Treg cell snATAC-seq and snRNA-seq. (C) Heatmap of chromatin accessibility for detected genomic regions for each lung pseudobulk sample. Read counts for peaks were aggregated by sample. The quantile of sum of read counts were calculated, and peaks falling between 20th and 90th percentile of all pseudobulk samples were kept. The peak counts were normalized by DESeq2 median of ratios method. The top 5000 most variable peaks were selected and visualized by Z-scores in heatmap. (D) Heatmap of chromatin accessibility for detected genomic regions for various cell activation states in the lung pseudobulk sample. Read counts for peaks were aggregated by cell state. The quantile of sum of read counts were calculated, and peaks falling between 20th and 90th percentile of all pseudobulk samples were kept. The peak counts were normalized by DESeq2 median of ratios method. The top 5000 most variable peaks were selected and visualized by Z-scores in heatmap. (E) Heatmap of enriched motifs for lung C1 resting, C2 primed and C4/C8 activated states, with cutoffs adjusted p-value < 0.05 and average difference > 1. The average motif activity score in each cell state was calculated and the z-score normalization was applied across cell states for visualization in the heatmap. (F) Correlation plot with regression showing similar DEGs from both lung and spleen snRNA-seq. (G) Volcano plots of lung and spleen snRNA-seq differentially expressed genes (activated states (C4, C8) versus resting state (C1)) with bZIP genes labeled in blue.

The chromatin accessibility profiles revealed a comprehensive list of accessible loci in the resting state of lung Treg cells (Figure 3D), which did not appear in the resting state of spleen Treg, due to failure to pass the same cut-off thresholds (Figure 2I). Regardless, splenic Treg cells in the *Ms4a4b*^+^ primed state shared some common accessible loci with those in the resting state in lung. The *Ass1^+^* and *Itgb7^+^* activated Treg states had accessible loci that were inaccessible in the *Satb1^+^* resting and *Ms4a4b^+^* primed states, and vice versa. The list of accessible/inaccessible loci shown in the lung cell state heatmap is summarized in Table S12.

Interestingly, significantly enriched motifs for both activated states in the spleen (Figure 2I) and lung (Figure 3E) were very similar, including, but not limited to, BATF, BATF3, BACH1/2, and the other bZIP TF motifs as well as the SMAD2/3 complex motif. Of note, the RUNX1 motif appeared to be accessible specifically in lung *Ms4a4b*^+^ primed Treg cells as well as in lung Treg *Ass1^+^* and *Itgb7^+^* activated states, whereas it was only enriched in spleen *Ass1^+^* and *Itgb7^+^* activated Treg cells when loosening the stringency of the cut-off values (log2FC) (Figure S8). Therefore, it suggests that increasing RUNX1 motif accessibility may play a critical role in the transition of primed Treg cells from the resting state to activated states in the lung. Although many enriched motifs in resting Treg cells from spleen did not reach the significance cutoff threshold, lung Treg cells did yield a short list of significantly enriched motifs based on snATAC-seq data. This list is comprised of TEADs, which corresponds to gene regulators important in homeostasis, as well as SOX and NRF family motifs, which correspond to gene regulators that are important in T cell stemness, maintenance, differentiation and metabolism ((Jiang et al., 2024), Moskowitz et al. (2017). Moreover, lung snRNA-seq data correlated with the spleen snRNA-seq data (Figure 3F), and the expression levels of bZIP TF family genes were similarly modulated between the resting and activated Treg cells (Figure 3G).

### IL-2 activated Treg cells demonstrate molecular phenotypes of T-helper-like cells

In attempt to correlate Treg activation with T-helper-like characteristics, we identified the co-expression and co-accessibility of activation marker genes with T-helper signature genes. Our spleen and lung snRNA-seq data suggest that IL-2 moderately upregulates expression of Th1 marker genes (*Ifng*, *Stat1*, *Tbx21*, *Cxcr3*) in *Ass1+* activated Treg cells, but downregulates the same genes in *Itgb7+* activated Treg cells. Expression of Th2 marker genes (*Gata3*, *Ccr4*, *Il4*, *Il13*, *Stat6*, *Irf4*) is upregulated in both spleen and lung *Ass1+* and *Itgb7+* activated Treg cells. Additionally, T-follicular helper (Tfh) marker genes (*Cxcr5*, *Bcl6*, *Pdcd*) appear to be expressed in the resting/primed and activated Treg cells, whereas expression of Th17 marker genes (*Rorc*, various *Il17 forms*, *Il1b*, *Il23*) is most upregulated in the *Satb1+* resting Treg cells (Figures 4A and 4B). These observations suggest that activated Treg cells show intrinsic demonstrable Th1, Th2 and partial Tfh molecular phenotypes. Similarly, in spleen and lung snATAC-seq, the IL-2 induced *Ass1+* and *Itgb7+* activated Treg cells show upregulated predicted gene activity for Th1 marker genes (*Ifng*, *Stat1*, *Tbx21*, *Cxcr3*), Th2 marker genes (*Gata3*, *Ccr4*, *Il4*, *Il13*, *Stat6*, *Irf4*), and Tfh marker genes (*Cxcr5*, *Bcl6*, *Pdcd*), but inconsistent activity for Th17 marker genes (*Rorc*, various *Il17 forms*, *Il1b*, *Il23*) (Figures 4C and 4D).

**Figure 4.**
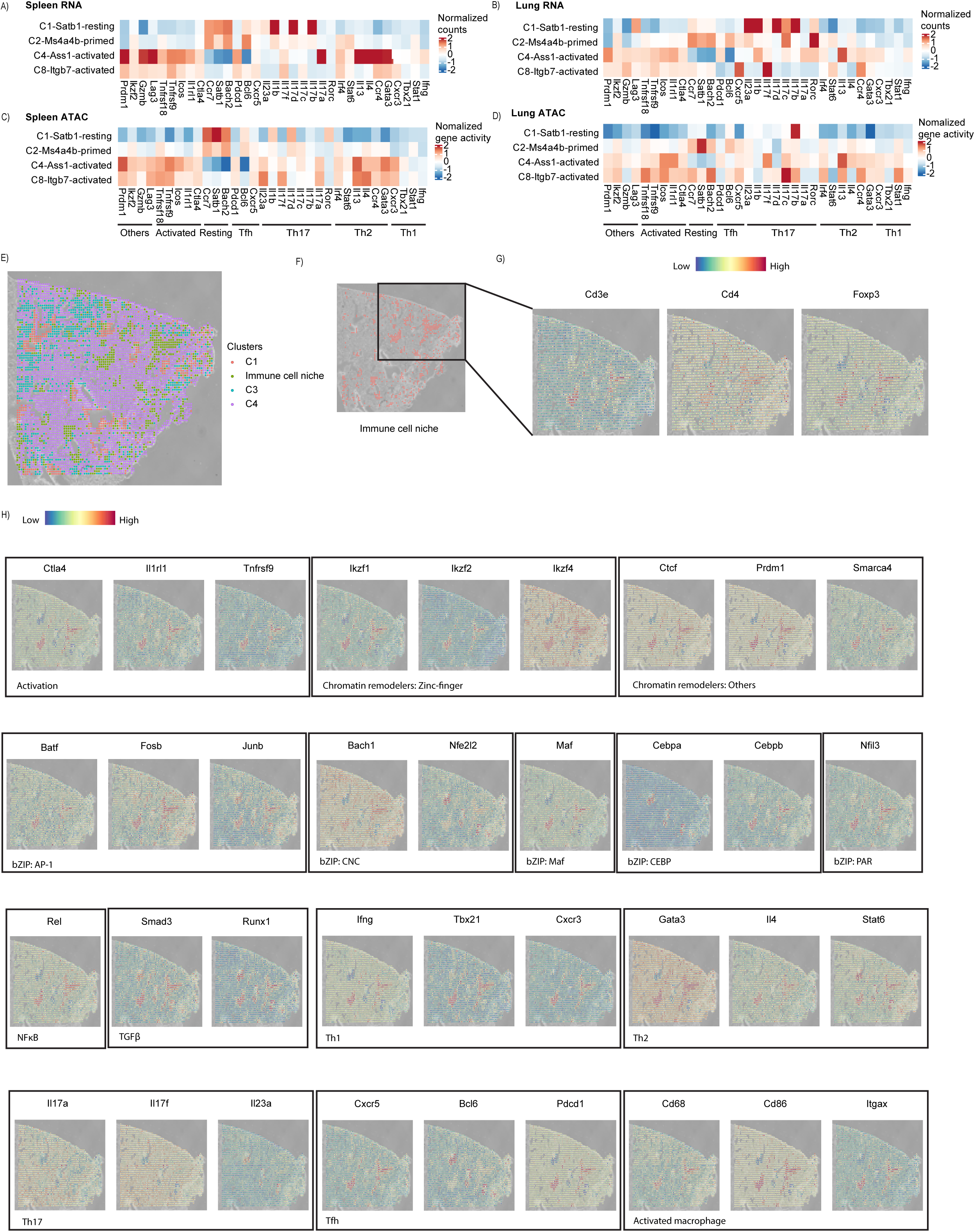
T-helper 1/2 cell features in IL-2 stimulated Treg cells. Heatmaps of aggregated normalized gene expressions, in counts, of T-helper cell signature genes from single-nuclei RNA-seq of spleen (A) and lung (B) Treg cells. Heatmaps of aggregated normalized chromatin accessibility, in gene activities, of T-helper cell signature genes from single-nuclei ATAC-seq of spleen (C) and lung (D) Treg cells. (E) Near single-cell resolution (20 µm) DBiT-seq based spatial ATAC-seq analysis at low clustering resolution identified an immune cell niche based on marker gene accessibilities. (F) The immune cell niche was emphasized for easy identification, with colocalizing (G) *Cd3e*, *Cd4* and *Foxp3* gene chromatin accessibilities, as well as (H) activation, chromatin remodeling, bZIP TF, and T-helper cell marker genes, in addition to activated macrophage markers.

To build confidence in the observed T-helper-like molecular phenotypes in Treg cells, we employed spatial ATAC-seq at single-cell resolution (20 µm), using a derivative method of DBiT-Seq platform (Liu et al., 2020, Deng et al., 2022), to assess IL-2M-treated mouse lung tissues. According to literature, lung Treg cells express higher levels of ST2, also known as a tissue Treg marker that is encoded by *Il1rl1*, than lymphoid Treg cells (Delacher et al., 2020b). Spatial ATAC-seq analysis on PBS control and mIL-2M treated mouse lung tissues revealed four clusters of cell niches at low clustering resolution (Figure 4E), including an immune cell niche (Figure 4F), which was absent in the control (data not shown) and further refined based on annotations of differential signature genes with high gene scores in *Ptprc* (immune cells), *Cd3d* (T cells), *Cd40lg* (B cells), *Cd68* (macrophages) and other immune cell markers (*Ccr2*, *Tnf*, *Ccr8* and *Ccl5*) (Table S13). Importantly, spatial overlap of the immune cell niche with marker genes *Cd3e* (pan-T cells), *Cd4* (CD4^+^ T cells) and *Foxp3* (Treg cells) confirms Treg presence (Figures 4G). Such “Y-shaped” Treg-rich regions (and associated spots to their left) also spatially overlap with activation markers (*Ctla4*, *Il1rl1*, *Tnfrsf9*), chromatin remodelers (*Ctcf*, *Prdm1*, *Smarca4*) and zinc finger TFs (*Ikzf1*, *Ikzf2*, *Ikzf4*), various bZIP TF genes (*Batf*, *Fosb*, *Junb*, *Bach1*, *Nfe2l2*, *Maf*, *Cebpa*, *Cebpb*, *Nfil3*) and other non-bZIP TFs, including *Rel* from NFκB signaling, *Smad3* from TGFβ signaling, and *Runx1,* a *Foxp3* transcriptional activator (Bruno et al., 2009), which were also identified as important regulators of Treg activation in our bulk and single-nucleus multiomics assays (Figures 4H and S11). Such activated Treg cells with enriched bZIP TFs showed T-helper-like molecular phenotypes, including Th1 marker genes (*Ifng*, *Tbx21*, *Cxcr3*), Th2 marker genes (*Gata3*, *Il4*, *Stat6*), and Tfh marker genes (*Cxcr5*, *Bcl6*, *Pdcd1*) but not Th17 marker genes (*Il17a*, *Il17b, Il23a*) (Figure 4H). Interestingly, the Treg cells colocalize with activated alveolar macrophages (*Cd68*, *Cd86, Itgax)*, suggesting potential involvement of cell-to-cell communication between activated Tregs and activated macrophages in non-lymphoid tissue after mIL-2M stimulation.

### Inhibition of cell activation upon bZIP TF gene perturbation

To experimentally test the impacts of bZIP TFs on Treg activation, Treg cells after perturbation (single or dual gene knockdown) were labeled by antibodies and quantified for live ST2^+^/4-1BB^+^ (“activated”) and ST2^-^/4-1BB^-^ (“resting”) states by flow cytometric analysis. Interestingly, there was a negatively correlated linear trend showing increased resting Treg cell numbers with decreased activated Treg cell numbers upon single gene perturbation (Figure 5A). Overall, maybe due to expression mainly in the resting state, *Bach* family genes appear to have greater impacts on Treg activation than *Batf* family genes (BACH2>BACH1=BATF1>BATF3), whereas multiple components of JUN and FOS related AP-1 heterodimers showed different levels of linear correlation between loss of ST2^+^/4-1BB^+^ Tregs and gain of ST2^-^/4-1BB^-^ Tregs upon knocking out individual genes. The effect of AP-1 heterodimer knockouts demonstrated more synergistic inhibition than any single-gene knockouts (Figure S12A), with an observed in the impactful trend of *Junb* and then *Jund* followed by *Jun,* in parallel with a similar pattern of inhibitory strength from *Fosl2* to *Fosl1* and then *Fos* (Figure 5B). However, no additive effect of any bZIP TF dual knockout combinations involving *Batf* or *Bach* members was observed (Figures S12B and S12C). To examine the impact of bZIP TF perturbation in chromatin accessibility, selective gene knockouts of combinatorial bZIP TF members based on the motifs revealed by ATAC-seq were performed and assessed. Differential ATAC-seq peak analysis enumerated a total of 6543 significant (FDR ≤ 0.05) downregulated differential peaks upon *Fosl2* and *Jund* dual gene knockout, which were even greater than *Batf* single gene knockout at 5860 peaks (Table S14), supporting the observed augmented loss of ST2^+^/4-1BB^+^ Treg cells for the dual knockout. Furthermore, the impactful trend of *Fosl2* to *Fosl1* was supported by the total number of significant differential peaks (Table S14). Of note, the significant differential peaks upon knockouts of *Batf*, *Fosl1* and *Junb*, *Fosl1* and *Jund*, as well as *Fosl2* and *Jund* were detected at the gene loci that involved in chromatin remodeling (*Prdm1*, *Hdac4*), T cell activation (*Icos*, *Ctla4*, *Runx1*), bZIP TF regulation (*Bach2*, *Maf*) and IL-2 signaling (*Stat5b*). Such a linear correlation of gain versus loss between resting and activated Treg cell proportions suggests that bZIP TFs may behave as intrinsic factors, imposing chromatin DNA remodeling and/or transcriptional regulation during mIL-2M-induced Treg activation. Bulk ATAC-seq analysis also showed that either *Fosl2* and *Jund* dual knockdown or *Batf* single knockdown decreased accessible ATAC-seq peaks at the proximal regulatory region of *Il1rl1* (which encodes the ST2 protein), supporting the notion that BATF or FOSL2/JUND may regulate *Il1rl1* expression through the binding to different motifs at the *Il1rl1* promoter (Figure 5C). Furthermore, while *Batf* single knockdown slightly increased accessible ATAC-seq peaks at the proximal regulatory and gene body regions of the resting marker gene *Sell* (supporting the inhibition of Treg activation and the accumulation of resting Tregs), *Fosl2* and *Jund* dual knockdown decreased accessible ATAC-seq peaks at the same regions of the *Sell* locus (suggesting that *Fosl2* and *Jund* induced Treg activation have a more upstream role than *Sell* along the activation spectrum and potentially explains their efficacious effects in limiting Treg activation upon dual knockdown) (Figure 5D).

**Figure 5.**
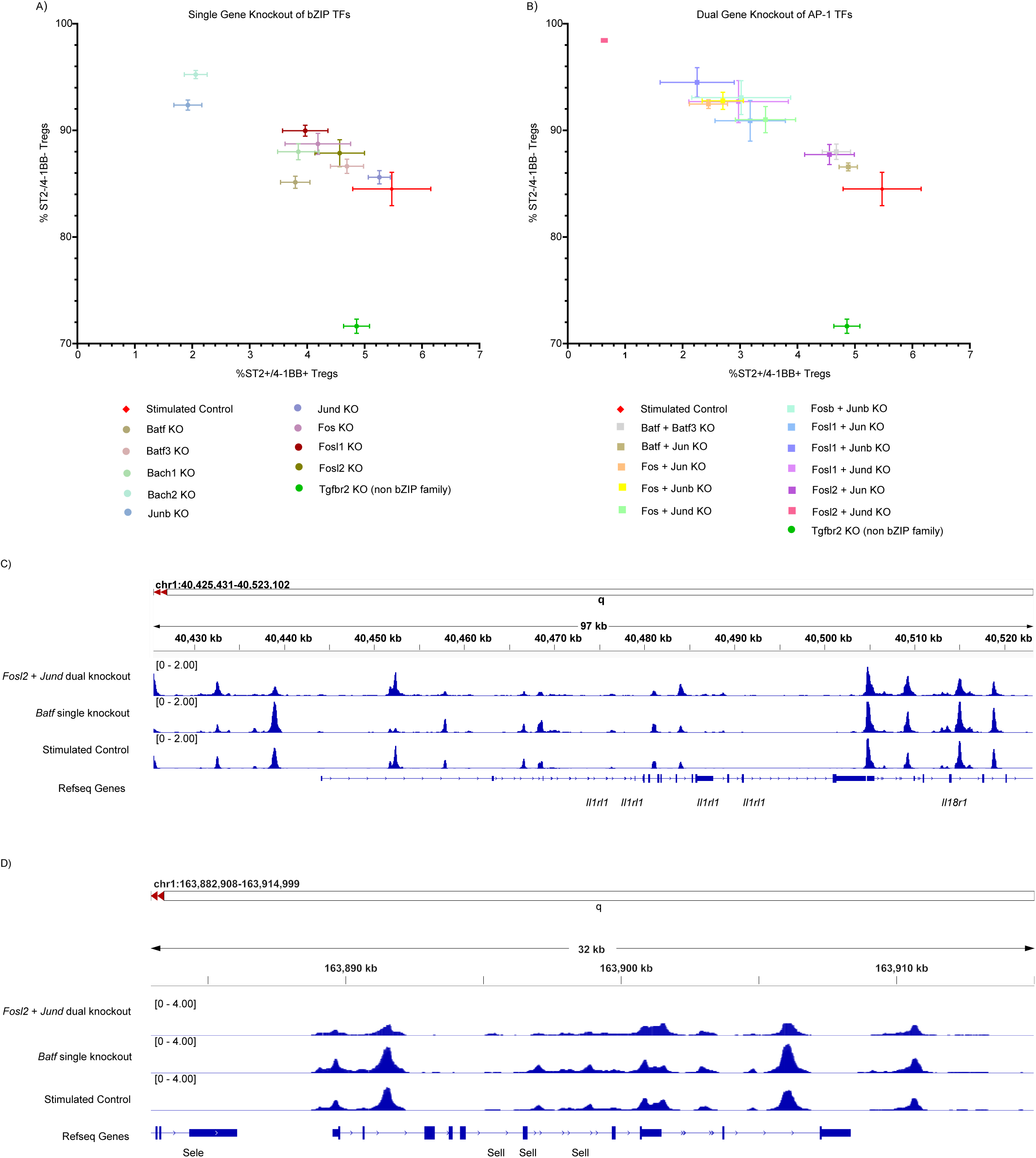
CRISPR knockout of basic leucine zipper transcription factors confirms their contributory role in Treg cell activation. (A) Percentage of live Treg cells that co-express ST2 and 4-1BB surface proteins compared to those that do not co-express them for each single gene knockout condition, where color filled circles represent means of technical triplicates and error bars represent standard deviations. (B) Percentage of live Treg cells that co-express ST2 and 4-1BB surface proteins compared to those that do not co-express them for each dual gene knockout condition, where color filled squares represent means of technical triplicates and error bars represent standard deviations. (C) Integrated Genomics Viewer (IGV) view of bulk ATAC-seq peaks comparing *Fosl2* and *Jund* dual and *Batf* single gene knockout with the control at the *Il1rl1* locus. (D) Integrated Genomics Viewer (IGV) view of bulk ATAC-seq peaks comparing *Fosl2* and *Jund* dual and *Batf* single gene knockout with the control at the *Sell* locus.

### IL-2 stimulation promotes BATF or BACH binding to bZIP DNA motifs

To understand how bZIP TF activities impact gene expression, CUT&RUN analysis on human primary Treg cells was performed to assess the physical binding of BATF (which plays a critical role in Treg activation as a transcriptional activator) and BACH1 (which is a tissue-agnostic ubiquitously-expressed BACH transcription repressor). Prior to this analysis, we checked whether the findings in mouse Treg cells would be translatable to human Treg biology by performing bulk ATAC-seq analysis on human primary Treg cells, which reproduced the bZIP TF motif panel identified and enriched in mouse IL-2-induced activated Treg cells (Table S15). Of note, ATAC-seq also identified CTCF motif as one of the most significantly enriched motifs in IL-2 treated human Treg cells, ranked by log P-value, followed by the motifs recognized by several ETS TFs. Subsequently, CUT&RUN analysis of histone marks revealed changes in annotated binding profiles after IL-2 stimulation, except H3K4me3 (Figure 6A). H3K27ac increased binding to enhancers after IL-2 stimulation, while H3K9me3, a repressive marker, decreased binding to intergenic and introns but increased binding to promoter and 5’UTR or exons.

**Figure 6.**
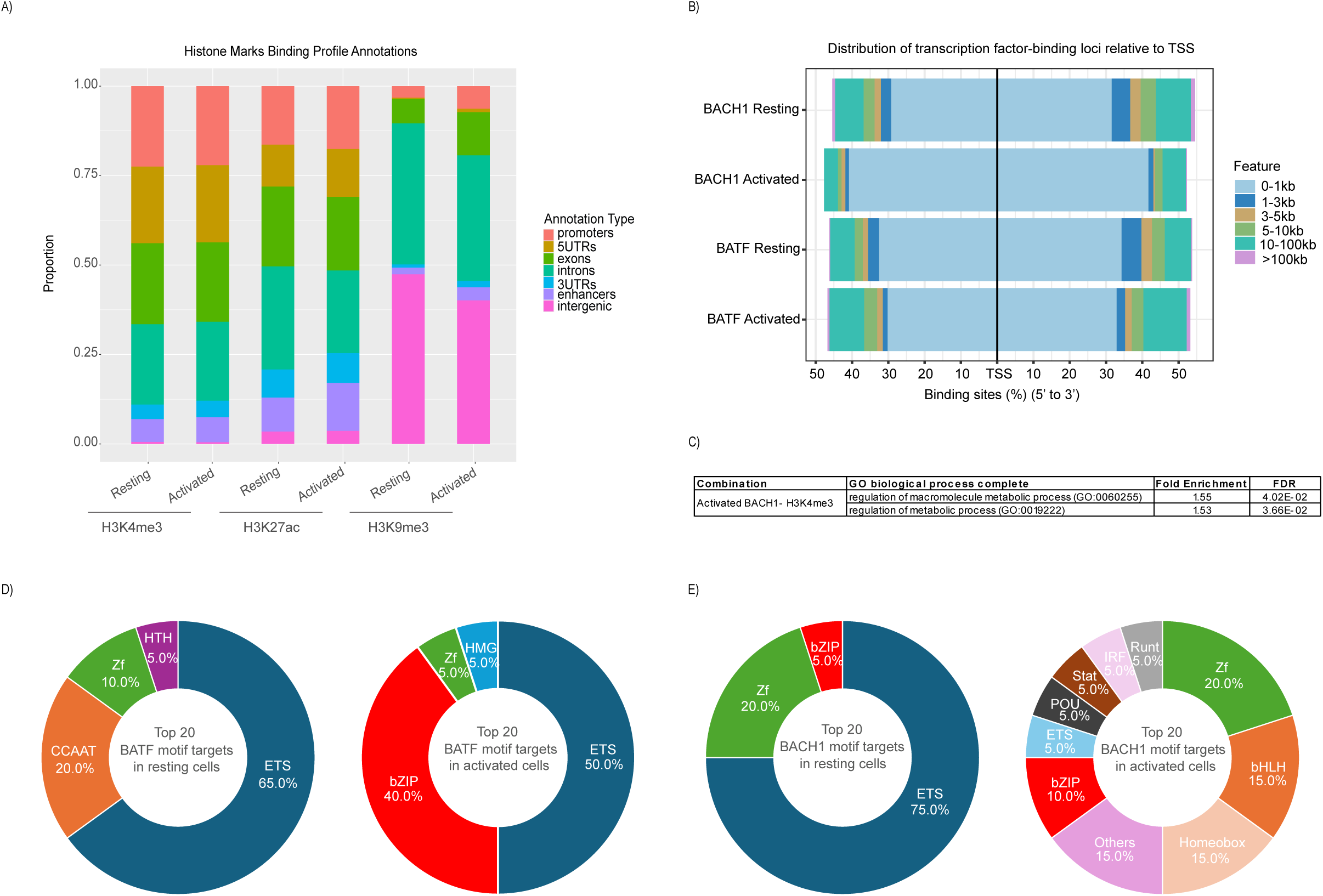
Dynamic chromatin remodeling increases bZIP binding targets. (A) Binding profile annotations of histone marks in human Treg cells with and without IL-2 stimulation by CUT&RUN assay. (B) Expansion of core promoter BACH1 and extension of distal BATF binding upon IL-2 stimulation. (C) Gene Ontology (GO) analysis based on the PANTHER database identified significant (FDR < 0.05, binomial distribution) metabolic process enrichment on overlapping BACH1 and promoter H3K4me3 peaks in activated Treg cells. Proportions of the top 20 enriched TF family binding targets of (D) BATF and (E) BACH1 in naïve and activated human primary Treg cells.

CUT&RUN analysis also demonstrated that both tested bZIP TFs (BATF and BACH1) were mainly bound to promoter regions with or without IL-2 stimulation. Intriguingly, the DNA binding profiles of both bZIP TFs were expanded upon IL-2 stimulation (Figures S13A and S13B). Specifically, while BATF binding extended from core promotors to distal regulatory regions relative to transcription start sites (TSS), BACH1 binding only expanded core promoter regions (Figure 6B). This expanded core promoter binding involved genes enriched in metabolic processes by GO analysis (Figure 6C), as well as genes specific for T cell specification (*CD3D*), T cell activation (*JUND*, *RUNX2*) and chromatin remodeling (*IKZF3*, *HIST1H3J*, *HIST2H2BE*) (Table S16). Furthermore, BATF and BACH1 bindings to bZIP annotated peaks were significantly increased post IL-2 activation (Figures 6D and 6E), with BACH1 mainly increasing the proportion of bound peaks at promoters, whereas BATF binding increases in more distal sequences upstream of the TSS. These observations suggest that BATF not only promotes target gene expression by binding proximally to the core promoters, but also regulates target gene activation via distal upstream regulatory elements, such as enhancers, by engaging in long-range cis-acting interactions, or insulators, by differentially regulating formation of TAD-domains (Figures S14A and S14B). The latter seems to be more likely, according to the observations that the BATF CUT&RUN peaks in activated Tregs were enriched with ETS and zinc-finger (ZF) motifs, which could be bound by CTCF, as well as high-mobility group (HMG) motifs, which are also known as a class of non-histone DNA regulators with functions in gene regulation (Ozturk et al., 2014), and the BATF CUT&RUN peaks overlapped with the enhancer mark H3K27ac did not significantly increased in activated Tregs. Upon further manual examination of these bZIP TF-bound loci in table S16, we found that ∼70% of chromatin DNA binding genes featured in the *Itgb7^+^* Tregs and ∼90% of TF genes profiled in the activated Treg cells were enriched both in BATF peaks in activated Treg cells and BACH1 peaks in non-stimulated Treg cells. In addition, several epigenomic regulator families, including *IKZFs*, *PRDMs*, and *HDACs*, increased DNA binding in both BATF activated and BACH1 unstimulated samples (Table S16). Taken together, these data imply that bZIP TFs, like BATF or BACH, may play a role in orchestrating chromatin accessibility profiles to establish a transcriptional expression program for inducing or maintaining Treg activation states after IL-2 stimulation. To further illustrate the functions of BATF and BACH1 target genes of upon IL-2 stimulation, we performed GO analysis for Biological Processes on genes whose promoters demonstrated binding for these bZIP TFs (Table S17). In line with our earlier results, BACH1 was classified as a transcriptional repressor and further identified as a regulator of metabolic processes. Also shown in Table S17, upon loosening the significance cutoff stringency, BACH1 was also identified as a positive regulator of T cell activation and Th1/2/fh cell differentiation, as well as B cell differentiation and activation. Simultaneously, it appears to be a negative regulator of macrophage differentiation and activation, suggesting promotion of the adaptive rather than innate immune system. Similarly, BATF target genes were enriched in cellular responses to IL-12 (which induces Th1 cell differentiation), dendritic cells (including macrophages), and regulation of cellular signaling and communication. As with BACH1, loosening the significance cutoff identified BATF target genes as enriched in positive regulation of Th1/2/fh cell differentiation and B cell activation and differentiation, but in negative regulation of macrophage activation.

### BATF or BACH perturbed primary Treg transcriptomes

To further illustrate the effects of BATF and BACH on T-helper-like characteristics, Treg cells were “inhibited from activation” by modulating candidate pro-activating bZIP TF genes’ expression via genome-editing studies, followed by scRNA-seq. Gene perturbation (knockdown) of *BATF* or *BACH1* in human primary Treg cells, followed by 10x Genomics scRNA-seq, confirmed their putative roles in “potential programming” of activated Treg cells with T-helper-like transcriptomic profiles. Specifically, *BATF* knockdown led to significantly decreasing the expression levels of Th1 marker genes (*CXCR3* and *IL12RB2*), Th2 marker genes (*GATA3* and *MAF*) and Tfh marker genes (*SLAMF6* and *CCR7*) compared to the unperturbed control (Figure 7A). *BACH1* knockdown resulted in decreased expression levels of Th1 marker genes (*NKG7* and *TNF*) and moderate decrease of Th2 marker genes (*MAF*). We note that human Treg cells with or without candidate bZIP TF gene perturbations barely detect appreciable levels of Th17 marker genes (*IL17*, *RORC*, *IL22*, *CCR6* and *IL23R*). Overall, this result also confirms the translatability of our mouse findings in human Tregs with respect to the role of bZIP TFs in cell activation and the demonstrable T-helper cell molecular phenotypes selectively regulated by BATF and BACH. Indeed, GSEA at the single cell level confirmed the loss Th1/2 phenotypes upon *BATF* or *BACH1* perturbation (Figures 7B and 7C).

**Figure 7.**
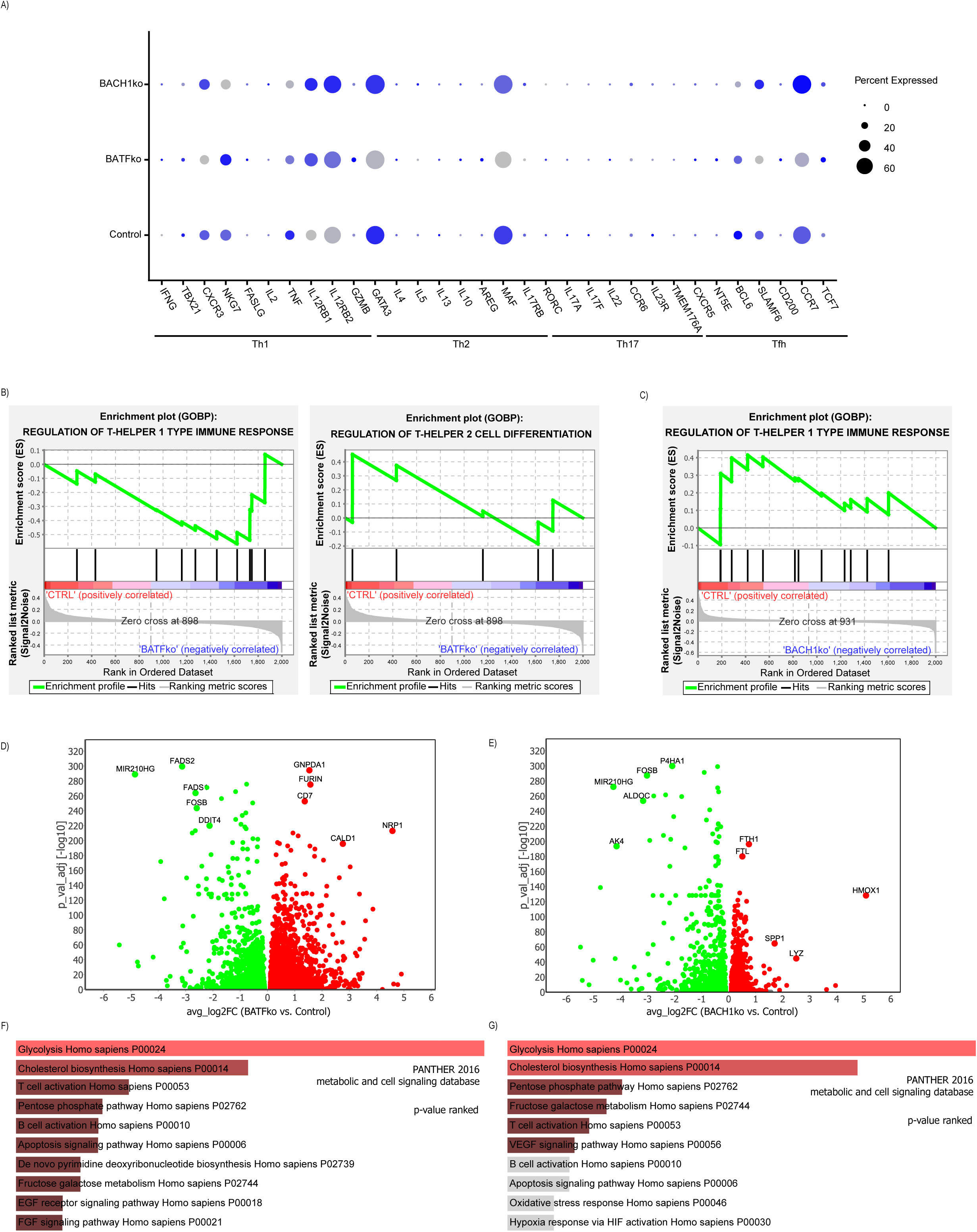
CRISPR knockdown of BATF and BACH basic leucine zipper transcription factors confirms their role in regulating T-helper cell features in stimulated human Treg cells. (A) Dot plot of pseudo-bulk T-helper cell signature gene expression levels after knockdown of BATF and BACH1 (plus control) by Perturb-seq. Single-cell level Gene Set Enrichment Analysis (GSEA) based on T-helper cell gene sets identified enrichment in T-helper cell 1 and 2 immune response and differentiation upon (B) BATF and (C) BACH1 knockdown. Perturb-seq pseudo-bulk differentially expressed genes (DEGs) upon (D) BATF and (E) BACH1 knockdown identified top 5 significantly upregulated (red) and downregulated (green) DEGs. EnrichR analysis using the PANTHER 2016 metabolic and cell signaling database of the downregulated genes upon (F) BATF and (G) BACH1 knockdown showed enrichment in metabolic processes.

To explore a more global view of cellular pathways upon *BATF* or *BACH1* perturbation, the respective pseudo-bulk DEGs of IL-2 stimulated versus unstimulated Treg cells were examined. *BATF* and *BACH1* perturbation downregulated some bZIP TF members (Figures 7D and 7E). GO pathway analysis of downregulated DEGs upon BATF perturbation corroborated loss of the T cell activation pathway, while BACH1 perturbation was validated by upregulation of HMOX1. Interestingly, similar highly significant metabolic genes were downregulated after either BATF or BACH1 perturbation, and GO analysis identified not only shared perturbed pathways in T-cell activation, apoptotic signaling, and metabolic processes, including glycolysis, cholesterol biosynthesis and pentose or fructose metabolism, but also distinct ones in EGF or FGF signaling and pyrimidine biosynthesis found in *BATF* knockouts and VEGF signaling or HIF activation specific for the *BACH1* KO Tregs, respectively (Figures 7F and 7G).

## Discussion

Here, we systematically characterized chromatin accessibility in mouse Treg cells activated by mIL-2M and confirmed that dynamic chromatin remodeling across Treg cell activation states was accompanied by changes in gene expression. Comparable datasets generated from bulk and single-nucleus multi-omics assays of resting and activated Treg cells allowed us to identify critical TFs that control Treg proliferation, differentiation and activation. This control appears to be mediated by increases in chromatin accessibility and binding to relevant motifs in spleen and lung, suggesting a comparable mechanism of IL-2-induced Treg activation functions in both lymphoid and non-lymphoid tissues. Our work illustrates a broad Treg activation regulome, whereas most major studies to date have focused on selected transcription factors or FOXP3-dependent regulation. Our results highlighted that bZIP TF-bound motifs were enriched in response to IL-2 stimulation, in addition to regulatory elements of the nuclear effectors for TGFβ and NFκB canonical pathways. Some of the identified bZIPs have been reported to regulate the epigenome of resting and activated T cells (Zhong et al., 2022), including BATF1, MAF, BACH2, JUNB, or FOS; the others, including BACH1, FOSL2, and JUND, are novel, or not well-defined, in Treg function or activation. The results of our gene perturbation experiments, followed by phenotypic, transcriptomic, and/or epigenomic analyses, on these bZIP TFs, confirmed their essential roles in IL-2-induced Treg activation in both mouse and human cells.

The bZIP TFs are exclusively eukaryotic proteins that bind to sequence-specific double-stranded DNA as homo- or heterodimers to either activate or repress transcription (Vinson et al., 2002). It has been reported that at least 53 unique genes (grouped into 23 families) of the human genome contain a bZIP binding motif, representing combinatorial complexity to form 2809 dimers, which enables a wide breadth of flexible, yet critical, transcriptional regulation. We demonstrated that combinatorial gene perturbation of *Fos* and *Jun* members, two typical bZIP TFs of the AP-1 family, suppressed cell activation more than single gene knockout of either *Fos* or *Jun* family member except for the specific combination of *Fosl2* and *Jun*. We also revealed that the impact of the AP-1 complex on ST2^+^/4-1BB^+^ Treg activation is mediated by its complex-forming partners: FOSL2 as the strongest, followed by FOSL1, and then FOS; similarly, JUNB or JUND as the strongest, followed by JUN. It has been shown that “emergent” sites, comprising of 16% of DNA binding motifs, are unique to TF heterodimerization, indicating that bZIP heterodimers exhibit more complex DNA-binding specificity landscapes than their individual counterparts (Rodríguez-Martínez et al., 2017). Our results concur with this finding for most of the AP-1 complex knockouts. Further Treg ATAC-seq assessment after gene perturbation of the most impactful AP-1 complexes, including FOSL2/JUND or FOSL1/JUNB or BATF alone, led to differential chromatin accessibility in the vicinity of chromatin remodeler (*Prdm1, Hdac4*), T cell activator (*Icos, Ctla4, Runx1*), bZIP TF regulator (*Bach2, Maf*) and IL-2 signaling factor (*Stat5b*) in addition to Treg cell-state markers, *Sell* and *Il1rl1*.

Consistent with previous findings in pan-T or Treg cells, we also identified that BATF is a critical transcriptional and epigenetic regulator in mIL2-M induced mouse and IL2-treated human Treg cells (Zhong et al. 2022, Itahashi et al. 2022). Expression of *Batf* and other TFs, but not chromatin remodelers, was robustly elevated in the *Ass1^+^* population of ST2^+^/4-1BB^+^ Tregs, aligning with an increase in chromatin accessibility in activated Tregs, suggesting that epigenetic modulation may be coordinated by cis-acting elements and/or topological events. In addition to physical interactions on core promoters or long-range regulation potentially via enhancers, BATF has been reported to initiate gene transcription in developing effector CD4^+^ T cells or control Ets-1 regulated recruitment of the architectural factor CTCF in order to promote chromatin looping associated with Th17 or Tfh lineage-specific gene transcription (Pham et al., 2019). Our CUT&RUN results demonstrated that BATF peaks are enriched with ETS, HMG and CTCF sites, as well as the expected bZIP TF motifs, suggesting a BATF-initiated mechanism of transcription activation via TAD reorganization or chromatin looping may be implicated in the initiation of Treg cell activation. However, we do not underestimate or totally rule-out cis-acting regulation via enhancers, because the typical histone mark for enhancers, H3K27ac, proportionally increased in Treg cells once stimulated by IL-2 in our study. In fact, a forward genetic screen revealed the mechanisms of enhancer-mediated *IL-2* gene regulation in Jurkat or primary T cells, reporting the involvement of several bZIP TFs (BATF3, JUN, JUND) (Pacalin et al., 2024). The CUT&RUN data also showed that, upon IL-2 treatment, BATF acts directly on multiple chromatin remodelers or histone modifiers, including PRDMs, HDACs, and IKZFs. This suggests that fine-tuning of chromatin remodeling by these factors may be sustained, but is predominantly crucial along the duration of Treg activation. Indeed, we observed that the motif activity and/or chromatin accessibility was slightly altered in the *Itgb7^+^* subpopulation of ST2+/4-1BB+ activated Tregs, whose signature genes list was enriched for chromatin DNA binding factors. Our results reveal that Treg heterogeneity during activation is dynamic and regulated by distinct epigenomic mechanisms depending on cell transitions or states.

Our analyses of the mouse single-nucleus RNA-seq and ATAC-seq data have provided evidence to elucidate the reprogramming and differentiation of T-helper cell-like molecular phenotypes in IL-2 induced Treg cells via bZIP TF regulation. In addition to bZIP TFs, epigenomic regulators, and activation markers, IL-2 induced Treg subsets up-regulated expression and increased chromatin accessibility of several Th1, Th2 and Tfh cell signature genes compared to the resting and primed Treg subsets. This was confirmed further by our spatial ATAC-seq data, in which we observed spatial colocalization of accessible *Foxp3* and Treg activation markers along with Th1, Th2 and Tfh cell signature genes and bZIP TFs. This demonstrated that activated Treg cells with T-helper associated molecular characteristics is consistent with previous studies, which showed that Treg cells can phenotypically mirror T-helper cell subsets in health and disease (Trujillo-Ochoa et al., 2023). In our study, human Treg cells expressed Th1, Th2 and Tfh cell signature genes when cultured in IL-2. Gene signatures of Th1, Th2 and Tfh were selectively downregulated in *BATF* KO or *BACH1* KO cells, confirming the critical roles of bZIP TFs in selectively mediating the expression of effector T-helper cell feature genes in Treg cells. Our orthogonal results from mouse and human data further support the cross-species translatability of our findings. Indeed, bZIP TFs are highly conserved between mouse and human; for example, mouse and human BATF and BACH1 share 90% and 80.3% in sequence homology, respectively *(Mestas and Hughes, 2004, Hu et al., 2024).* Although it is not well-known why Treg cells reprogram to sustain Th-like molecular phenotypes, in agreement with colocalization of activated macrophage marker genes in our spatial ATAC-seq data, a recent study revealed that the Treg cells which polarize into Th1 feature post stimulation of tumor-associated macrophage (TAM) secreting platelet factor 4 (PF4), possibly attenuating anti-tumor immunity and resulting in enhancement of tumor growth (Kuratani et al., 2024). Nevertheless, it also noted that Th17 cell signature genes were not detected in either of our mouse or human datasets. This may be because the lineage-specifying TF for Th17 cells (e.g. RORγt) is more abundantly expressed in gut tissues regarding to the presence of microbial and food antigens. However, our data from mouse spleen and lung Treg cells and human PBMC Treg cells reflect more like healthy homeostatic states, which did not sustain detectable frequencies of gut-associated or pathogen-induced Treg cells.

BACH proteins in the digestive tract have been demonstrated to directly or indirectly regulate inflammation and tumor angiogenesis via modulation of physiological processes, including differentiation of T cells and B cells, mitochondrial function and heme homeostasis, as well as pathogenic processes, such as responses to infections, autoimmune disorders and changes in metabolism (Song et al., 2023). *Bach1* is a tissue-ubiquitous master regulation of oxidative stress (Zhang et al., 2018), including the anti-oxidant pathway in pathogen-induced inflammatory gut Treg cells (Vaikunthanathan et al., 2023). It has been shown that Treg metabolism is important in Treg activation via epigenetic mechanisms (Lu et al., 2021). However, supporting evidence specifically involving *Bach1* in Treg cell metabolism and activation state has been lacking. Existing literature does describe the role of *Bach2*, the tissue specific form of *Bach* family members, in the gastrointestinal tract in dampening TCR signaling via limiting chromatin accessibility (Sidwell et al.). *Bach2* also has a well-described role in B and plasma cell differentiation, a known role in some neuronal cell types and emerging roles in macrophage and T cell transcriptional activity at super-enhancers (Richer et al., 2016). It acts as a transcriptional activator or repressor in a context-dependent manner. Our data indicate that, in IL-2-treated Treg cells, *Bach1* and *Bach2* knockdown, either single or dual gene, decreased cell activation, suggesting that both are important in promoting Treg activation although the impact imposed by *Bach2* was more significant. It has been proposed that, given its predominant expression in the naïve state, *Bach2* stabilizes Treg development (before *Foxp3* induction) and promotes central Treg survival and differentiation, but can also repress genes associated with effector Treg differentiation from naive Treg cells *(Yang et al., 2019, Grant et al., 2020)*. We hereby identified *Bach1* as a novel regulator of Treg activation that may involve multiple metabolic processes in Treg cells (Delacher et al., 2020a, Delacher et al., 2021). However, the role of *Bach1* in Treg development and differentiation remains worthy for further study, especially since we showed that disruption of BACH1 expression ameliorated the formation of ST2^+^/4-1BB^+^ activated Tregs and inhibited expression of selective Th1-like marker genes (e.g. *NKG7* or *TNF*). Nevertheless, we found that, similar to direct BATF regulation of *FOSB* or *MAF* expression, knockouts of BACH1 disrupted expression of some AP-1 complex genes like FOSB in human Treg cells, suggesting that there may be multiple lines of feedback regulation mechanisms among bZIP TF members, which also needs further investigations.

As the examples of bZIP TFs of interest in this study, BATF and BACH1 seemed to regulate key genes in a wide range of physiological processes, including inflammation and immune response, metabolism, cell proliferation and differentiation, and stress response, rendering them attractive targets to intervene upon in disease processes. Indeed, GO analysis of snRNA-seq DEGs from mouse *Ass1*^+^ and *Itgb7*^+^ activated Treg states using the Reactome 2022 database showed enrichment in metabolic pathways (data not shown). GO analysis on BACH1 CUT&RUN peaks confirmed some BACH1 target genes are involved in metabolic processes. Simultaneously, our Perturb-seq data showed that *BATF* or *BACH1* knockdown in Treg cells downregulates metabolic gene expression, and GO analysis of these downregulated genes confirmed the contribution of BATF and BACH1 to metabolic processes, in supportive of a positive regulatory role of these two bZIP TFs in promoting multiple metabolic pathways during IL-2 induced Treg activation. In addition to cholesterol metabolism, similar to BATF, BACH1 enabled the promotion of glycolysis, which aligns well with the published role of BACH1 in favoring cytoplasmic glycolysis over mitochondrial oxidative phosphorylation in a tumor microenvironment (Hu et al., 2024). An established role of BACH1 in cancer cells is to preferentially employ glycolysis instead of oxidative phosphorylation despite abundant oxygen levels, and this phenomenon is referred to as “aerobic glycolysis.” BACH1 regulates aerobic glycolysis to promote tumor growth, progression and metastasis in various cancers (including breast, lung, liver, glia), by inhibiting expression of mitochondrial electron transport chain genes and inducing expression of glycolytic enzymes. Accordingly, inhibiting BACH1 or its downstream effectors is a potentially viable anti-cancer therapeutic strategy. Indeed, we showed that BACH1 motif activity was increased in IL-2 stimulated activated Treg states that have demonstrated high suppressive Treg function, resulting in reduced anti-cancer immunity by suppressing effector T cell functions. Consequently, inhibiting BACH1 motif activity, or BACH1 binding sites and its target genes, could ideally either maintain Treg cells in the resting/naïve state or reverse activated Treg cells back to the resting/naïve state, both of which would result in lowering suppressive Treg cell function, thus increasing anti-cancer immunity by promoting effector T cell functions. Our results support the translatability of findings across mouse and human, and confirm the validity of the proposed BACH1 inhibition therapeutic strategy. Conversely, it has been previously shown that IL-2 stimulated mouse pan-Treg cells exhibit increased oxidative phosphorylation. This discrepancy may be a result of the forms of IL-2 used and the resultant target cell specificity. Nevertheless, further studies are needed to prove the hypothesis that BACH1 inhibition can promote Treg suppressive function in anti-cancer treatment.

Most prior literature has described Treg activation within the bounds of short-term stimulation (in hours). However, one previous study examined bulk transcriptome and chromatin accessibility profiles of prolonged CD25 stimulation (in days) and Treg activation using a modified IL-2 agonist and concluded that sustained stimulation induced upregulation of genes involved in cell proliferation and metabolic activity, and increased chromatin accessibility in STAT5 and CTCF motifs in the early and late phases of activation, respectively (Moro et al., 2022). In general, our results not only concur with these findings but also emphasize the enrichment of bZIP TF motifs in activated Treg cells. The significance of Treg cell activation state transitions may be relevant in a wide variety of homeostasis or disease indications. Specifically, epigenetic modulation of Treg activation states has been applied to and researched in viral infections, autoimmune- or tumor- induced inflammation, and even CAR-T cell therapy (Yamamoto-Taguchi et al., 2013, Mijnheer et al., 2021, Weber et al., 2021). Human T-cell leukemia virus type 1 (HTLV-1), for instance, encodes a bZIP gene that is constitutively expressed in infected cells, which results in increased Treg cell number by inducing *Foxp3* gene expression (Yamamoto-Taguchi et al., 2013).

In addition to the bZIP TF motifs, we identified other classes of motifs recognized by various TF families with important regulatory roles, including TBX, RUNX, IRF, PPAR, zinc-finger (e.g. SP, KLF, IKAROS), NFAT, ETS/ETV. Among these, KLF motifs, including KLF1, KLF3, KLF6 and KLF12, have been reported to function mainly in tissue homeostasis and metabolism (Ilsley et al., 2017, Pollak et al., 2018). TBX motifs, including TBX1, TBX15, TBX18 and TBX21, are known to regulate developmental processes, such as cell fate determination and cell differentiation (Wilson and Conlon, 2002). *Tbx1* has been shown to concentrate Treg cells into “immune hubs” organized by activated lymphatic endothelial cells during tissue repair in post-myocardial infarction heart (Wang et al., 2023). The involvement of TBX21 suggests T-BET programming plays a role in Treg cell activation towards T-helper 1 cell characteristics. In addition, RUNX1 is known to induce accessible chromatin and plays a key role in T cell activation (Korinfskaya et al., 2021). RUNX1 also induces *Foxp3* expression and interacts with specific regulators of T-helper cell subsets, including T-BET (Th1), GATA3 (Th2) and RORγt (Th17). The complex interplay of so many different TFs suggests the need for relatively high-dimensional molecular perturbation approaches to systematically investigate the regulatory mechanisms uncovered in the current studies.

In this study, we provide evidence that the bZIP superfamily of TFs serves as a critical class of transcriptional/epigenomic regulators for IL-2-induced Treg cell activation. However, in some bZIP TF cases, gene expression was not always directly proportional to or reflective of motif accessibility. For example, while bZIP TF motifs were enriched in activated Treg cells, gene expression levels of the corresponding bZIP TFs could be increased or decreased. This makes sense, given that some bZIP TFs are transcriptional activators, while the others are transcriptional repressors or could be bipolar (e.g. BACH1, FOSL2) although follow-up studies are needed to elucidate the mechanisms of action. Overall, our findings provide interesting directions for future studies to further characterize Treg epigenomic heterogeneity or interrogate epigenomic or transcriptional regulation during Treg state transition upon cytokine stimulation or disease conditions.

## Acknowledgments

We are thankful for Amgen *in vivo* Pharmacology group to provide the assistances of setting up mouse experiments and Discovery Protein Sciences group to kindly provide the study reagents. We also thank ASF flow cytometry core for the assistance of sorting Treg cells for the downstream assays and Discovery Genomics NGS/SC group for the help of all library sequencing. Lastly, we would like to give our sincere thanks to Hyewon Phee, Sue Sohn, Weiwen Deng, Daniel Ellwanger, Menno Van Lookeren Campagne, and Ryan Potts for the inputs on the proposal or the manuscript.

## Author Contributions

J.T. designed and executed the experiments. J.T. generated bulk multi-omics data. J.T. and T.Y. generated single-nuclei multi-omics data. J.T., X.Liu, M.Kanke, and I.E. analyzed data. J.T. and M. Killian. performed flow cytometry experiments. J.T. and A.S. extracted mouse tissue or performed Treg cell isolation. C.M.L., S.W., and A.C. supervised the study. J.L., D.L., W.L., C.Y.K., X.L., S.W., S.M., and A.C. provided inputs on experimental design, data analysis, or manuscript preparation. CM.L. initiated and finalized the research proposal. J.T. and CM.L. interpreted the data as well as drafted and finalized the manuscript.

## STAR Methods

### Key resources table

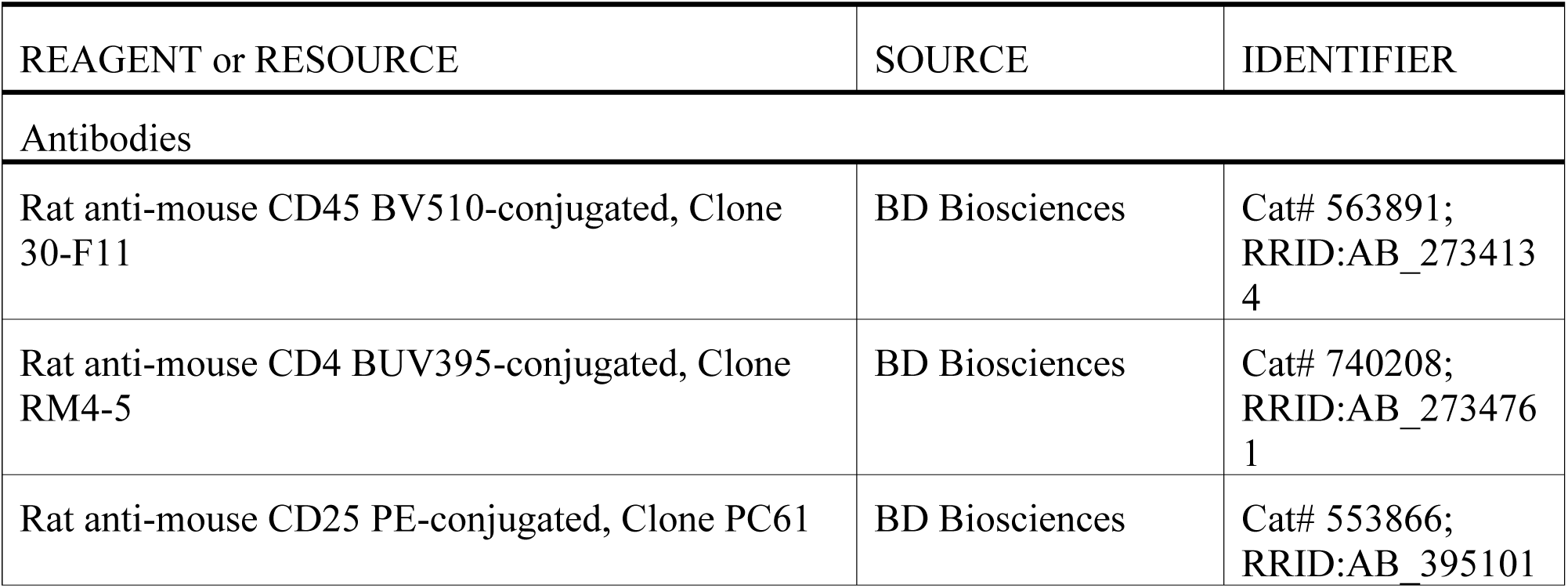
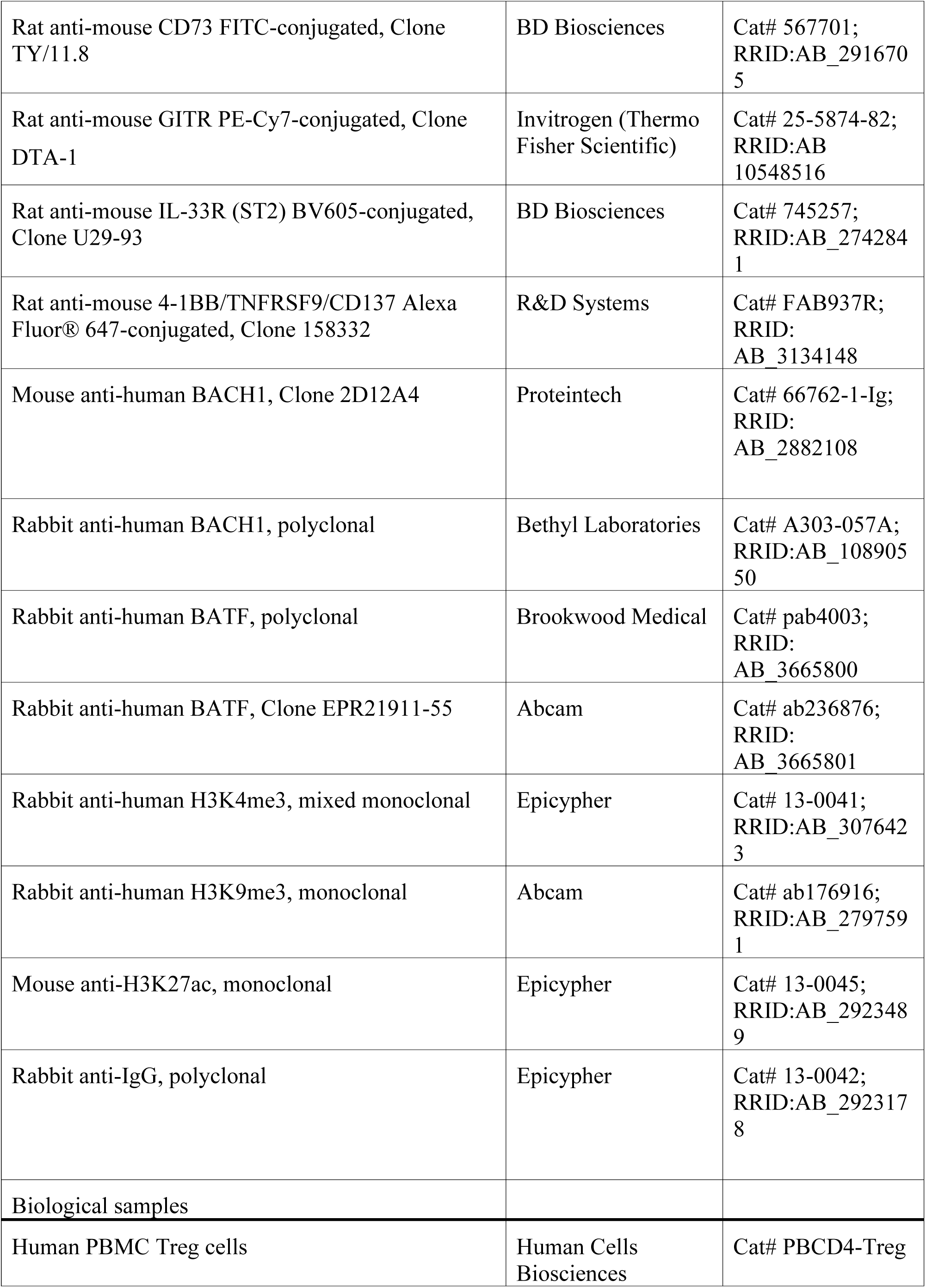
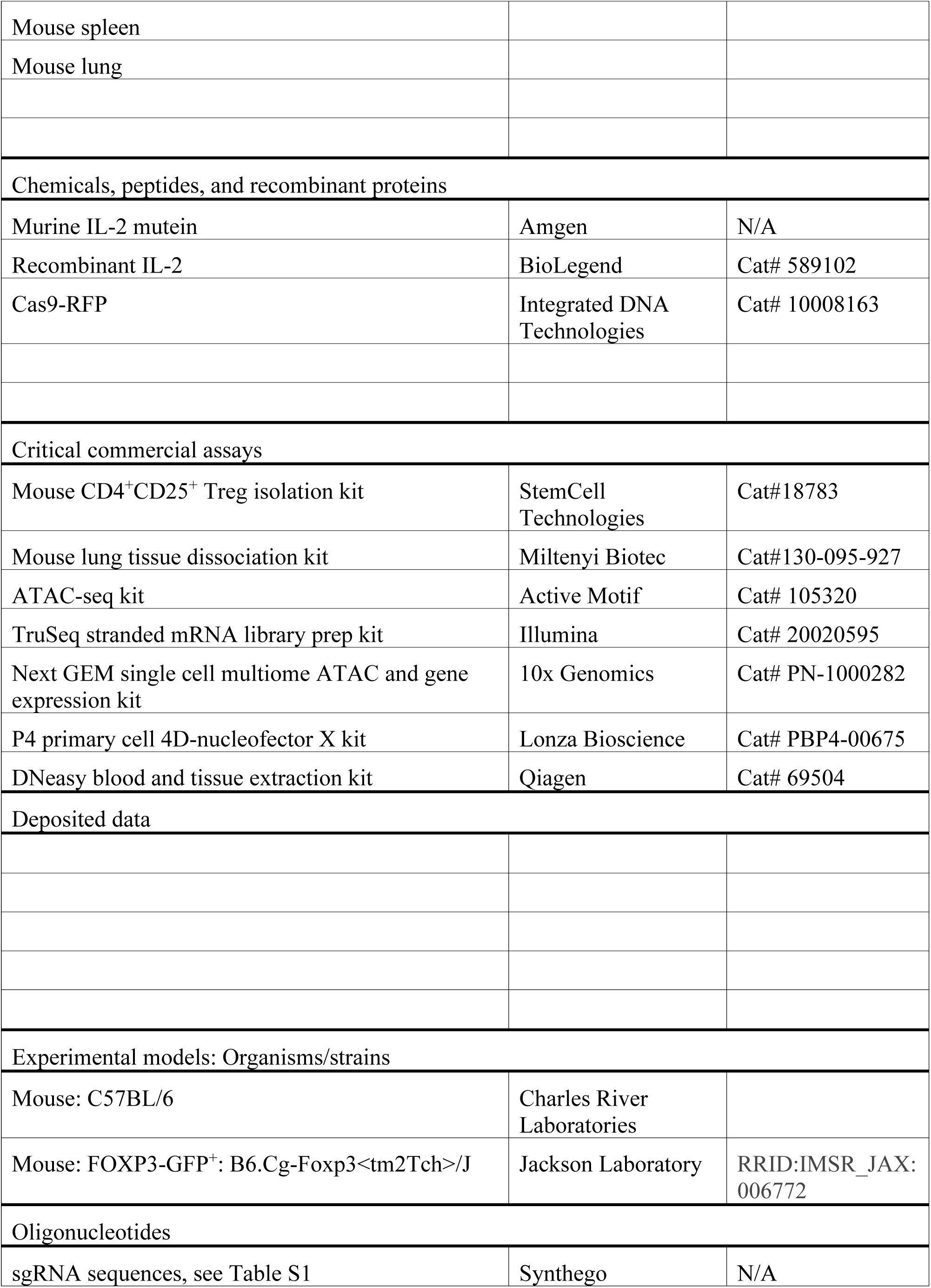
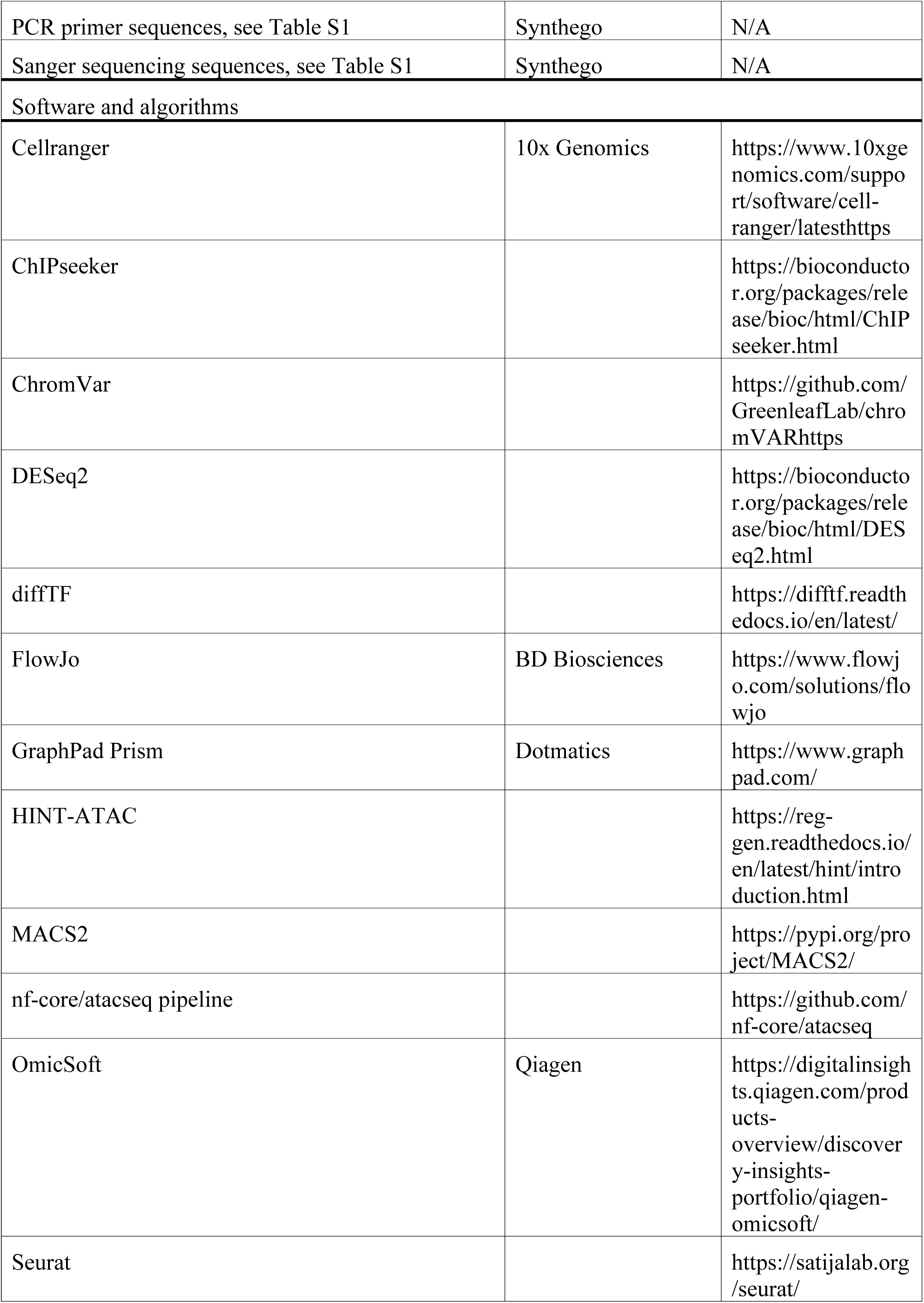
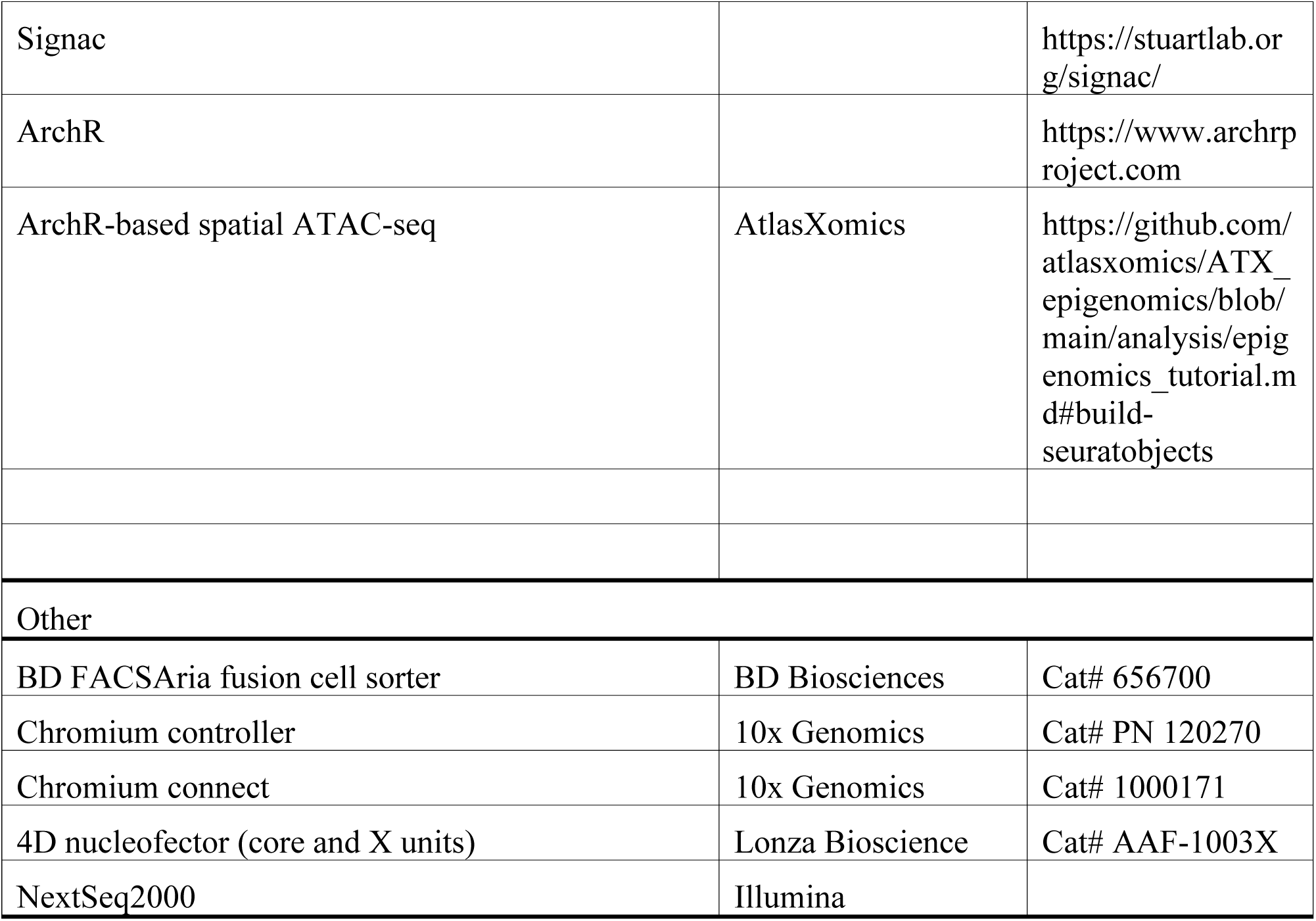

| REAGENT or RESOURCE | SOURCE | IDENTIFIER |
| --- | --- | --- |
| Antibodies |  |  |
| Rat anti-mouse CD45 BV510-conjugated, Clone 30-F11 | BD Biosciences | Cat# 563891; RRID:AB_2734134 |
| Rat anti-mouse CD4 BUV395-conjugated, Clone RM4-5 | BD Biosciences | Cat# 740208; RRID:AB_2734761 |
| Rat anti-mouse CD25 PE-conjugated, Clone PC61 | BD Biosciences | Cat# 553866; RRID:AB_395101 |
| Rat anti-mouse CD73 FITC-conjugated, Clone TY/11.8 | BD Biosciences | Cat# 567701;<br>RRID:AB_2916705 |
| Rat anti-mouse GITR PE-Cy7-conjugated, Clone DTA-1 | Invitrogen (Thermo Fisher Scientific) | Cat# 25-5874-82;<br>RRID:AB_10548516 |
| Rat anti-mouse IL-33R (ST2) BV605-conjugated, Clone U29-93 | BD Biosciences | Cat# 745257;<br>RRID:AB_2742841 |
| Rat anti-mouse 4-1BB/TNFRSF9/CD137 Alexa Fluor® 647-conjugated, Clone 158332 | R&D Systems | Cat# FAB937R;<br>RRID:AB_3134148 |
| Mouse anti-human BACH1, Clone 2D12A4 | Proteintech | Cat# 66762-1-Ig;<br>RRID:AB_2882108 |
| Rabbit anti-human BACH1, polyclonal | Bethyl Laboratories | Cat# A303-057A;<br>RRID:AB_10890550 |
| Rabbit anti-human BATF, polyclonal | Brookwood Medical | Cat# pab4003;<br>RRID:AB_3665800 |
| Rabbit anti-human BATF, Clone EPR21911-55 | Abcam | Cat# ab236876;<br>RRID:AB_3665801 |
| Rabbit anti-human H3K4me3, mixed monoclonal | Epcypher | Cat# 13-0041;<br>RRID:AB_3076423 |
| Rabbit anti-human H3K9me3, monoclonal | Abcam | Cat# ab176916;<br>RRID:AB_2797591 |
| Mouse anti-H3K27ac, monoclonal | Epcypher | Cat# 13-0045;<br>RRID:AB_2923489 |
| Rabbit anti-IgG, polyclonal | Epcypher | Cat# 13-0042;<br>RRID:AB_2923178 |
| Biological samples |  |  |
| Human PBMC Treg cells | Human Cells Biosciences | Cat# PBCD4-Treg |
| Mouse spleen |  |  |
| Mouse lung |  |  |
| Chemicals, peptides, and recombinant proteins |  |  |
| Murine IL-2 mutein | Amgen | N/A |
| Recombinant IL-2 | BioLegend | Cat# 589102 |
| Cas9-RFP | Integrated DNA Technologies | Cat# 10008163 |
| Critical commercial assays |  |  |
| Mouse CD4<sup>+</sup>CD25<sup>+</sup> Treg isolation kit | StemCell Technologies | Cat#18783 |
| Mouse lung tissue dissociation kit | Miltenyi Biotec | Cat#130-095-927 |
| ATAC-seq kit | Active Motif | Cat# 105320 |
| TruSeq stranded mRNA library prep kit | Illumina | Cat# 20020595 |
| Next GEM single cell multiome ATAC and gene expression kit | 10x Genomics | Cat# PN-1000282 |
| P4 primary cell 4D-nucleofector X kit | Lonza Bioscience | Cat# PBP4-00675 |
| DNeasy blood and tissue extraction kit | Qiagen | Cat# 69504 |
| Deposited data |  |  |
| Experimental models: Organisms/strains |  |  |
| Mouse: C57BL/6 | Charles River Laboratories |  |
| Mouse: FOXP3-GFP<sup>+</sup>: B6.Cg-Foxp3<sup>tm2Tch</sup>/J | Jackson Laboratory | RRID:IMSR_JAX:006772 |
| Oligonucleotides |  |  |
| sgRNA sequences, see Table S1 | Synthego | N/A |
| PCR primer sequences, see Table S1 | Synthego | N/A |
| Sanger sequencing sequences, see Table S1 | Synthego | N/A |
| Software and algorithms |  |  |
| Cellranger | 10x Genomics | https://www.10xgenomics.com/support/software/cellranger/latesthttps |
| ChIPseeker |  | https://bioconductor.org/packages/release/bioc/html/ChIPseeker.html |
| ChromVar |  | https://github.com/GreenleafLab/chromVARhttps |
| DESeq2 |  | https://bioconductor.org/packages/release/bioc/html/DESeq2.html |
| diffTF |  | https://difftf.readthedocs.io/en/latest/ |
| FlowJo | BD Biosciences | https://www.flowjo.com/solutions/flowjo |
| GraphPad Prism | Dotmatics | https://www.graphpad.com/ |
| HINT-ATAC |  | https://regen.readthedocs.io/en/latest/hint/introduction.html |
| MACS2 |  | https://pypi.org/project/MACS2/ |
| nf-core/atacseq pipeline |  | https://github.com/nf-core/atacseq |
| OmicSoft | Qiagen | https://digitalinsights.qiagen.com/products-overview/discovery-insights-portfolio/qiagen-omicsoft/ |
| Seurat |  | https://satijalab.org/seurat/ |
| Signac |  | https://stuartlab.org/signac/ |
| ArchR |  | https://www.archrproject.com |
| ArchR-based spatial ATAC-seq | AtlasXomics | https://github.com/atlasxomics/ATX_epigenomics/blob/main/analysis/epigenomics_tutorial.md#build-seuratobjects |
| Other |  |  |
| BD FACSAria fusion cell sorter | BD Biosciences | Cat# 656700 |
| Chromium controller | 10x Genomics | Cat# PN 120270 |
| Chromium connect | 10x Genomics | Cat# 1000171 |
| 4D nucleofector (core and X units) | Lonza Bioscience | Cat# AAF-1003X |
| NextSeq2000 | Illumina |  |

### Resource availability

#### Lead contact

Further information and requests for resources and reagents should be directed to and will be fulfilled by the lead contact, Chi-Ming Li.

#### Materials availability

This study did not generate new unique reagents.

#### Data and code availability

- The bulk RNA-seq, bulk ATAC-seq and 10x Multiome (single-nuclei ATAC-seq and single-nuclei RNA-seq) data reported in the study have been deposited at GEO: XXXX (to be updated) and are publicly available as of the date of publication. Accession numbers are listed in the key resources table. ICE analysis and flow cytometry data reported in this paper will be shared by the lead contact upon request.
- All original code has been deposited at Zenodo and is publicly available as of the date of publication. DOIs are listed in the key resources table
- Any additional information required to reanalyze the data reported in this paper is available from the lead contact upon request.

### Experimental model details

#### Animals

Adult female C57BL/6 and FOXP3-GFP^+^ mice (7-8 months old) were procured by Charles River Accelerator and Development Lab (CRADL) and The Jackson Laboratory, respectively. The animals were maintained by CRADL in 12 hour dark-light cycles with a set temperature of 22°C and humidity range between 30% to 70%. Mice were housed at no more than five mice per cage in ventilated racks (Innorack® IVC Mouse 3.5) with pre-filled ¼” Corn Cob in disposable cages from Innovive and fed the Standard Diet (#T.2920X. 10, Envigo). Housing enrichment included diamond half-twists and irradiated EnviroPaks as well as pre-filled acidified water bottle from Innovive.

### Method details

#### Murine IL-2 mutein treatment

Mice were administered with 10 ug of murine IL-2 mutein or phosphate buffered saline (PBS) for four days. For 10x Multiome assays, the agents were administered subcutaneously, while they were administered by i.p. injection for bulk multi-omics assays. For mouse CRISPR studies, 8 week-old female FOXP3-GFP^+^ mice were used.

#### Tissue dissociation and Treg cell enrichment

Murine spleens were mechanically dissociated into single cells and filtered through a 70 µm filter (#352350, Falcon). Filtered single splenocytes were resuspended in EasySep buffer (#20144, StemCell Technologies) and immunomagnetically enriched using magnetic beads by CD4 negative selection and CD25 positive selection to obtain CD4^+^CD25^+^ Treg cells (#18783, StemCell Technologies). Dissected mouse lungs were dissociated into single cells with a lung dissociation kit (#130-095-927, Miltenyi Biotec). In brief, single lobes of lung tissue were submerged in a buffered enzyme mix and dissociated with the gentleMACS Octo Dissociator with Heaters equipment (Miltenyi Biotec) under the 37C_m_LDK_1 program. Dissociated single cells were filtered through a 70 µm filter (#352350, Falcon), centrifuged and resuspended in EasySep buffer (#20144, StemCell Technologies), followed by immunomagnetic enrichment using magnetic beads to obtain CD4^+^CD25^+^ Treg cells as noted above (#18783, StemCell Technologies).

#### Flow cytometry and fluorescence-activated cell sorting of Treg cells

For benchmarking bulk multi-omics assays of resting and highly activated Treg cells, single splenocytes from mice injected with mIL-2M were subjected to red blood cell lysis (ACK RBC Lysis buffer pH 7.4 low endotoxin, Unity Lab Services, Thermo Fisher Scientific) prior to immunostaining and sorting. Red blood cell-lysed splenocytes were stained for dead cells using a near-infrared live/dead stain (#L34975, Invitrogen) in PBS for 30 minutes on ice in the dark. After two washes in PBS containing 2% FBS (#SH30071.03, Hyclone, GE Life Sciences), an antibody cocktail containing CD45-BV510 (#563891, BD Biosciences), CD4-BUV395 (#740208, BD Biosciences), CD25-PE (#553866, BD Biosciences), CD73-FITC (#567701, BD Biosciences), GITR-PE-Cy7 (#25-5874-82, Invitrogen), ST2-BV605 (#745257, BD Biosciences) and 4-1BB-AlexaFluor647 (#FAB937R, R&D Systems) were added to the cell suspension and incubated for 45 minutes on ice in the dark. After incubation, cells were washed 3 times with PBS + 2% FBS and subjected to BD FACSAria Fusion sorting. Cells double positive and double negative for ST2 and 4-1BB were collected in separate tubes containing Advanced RPMI1640 medium (#12633-012, Gibco) with 10% FBS (#SH30071.03, Hyclone, GE Life Sciences), kept at 4°C. Cells were pooled, such that each of two conditions (double positive and double negative) were in triplicates (total 6 samples) and pelleted by centrifugation, followed by bulk RNA-seq and ATAC-seq library preparation.

For CRISPR studies, red blood cell removed dissociated whole mice splenocytes were stained for dead cells using a near-infrared live/dead stain (#L34975, Invitrogen) in PBS, followed by resuspension in PBS containing 2% FBS (#SH30071.03, Hyclone, GE Life Sciences) for sorting of live FOXP3-GFP^+^ singlets that were collected in Advanced RPMI1640 medium (#12633-012, Gibco) with 10% FBS (#SH30071.03, Hyclone, GE Life Sciences), kept at 4°C. CRISPR knockout was conducted on the collected Treg cells, cultured and stimulated prior to flow cytometry assessment of ST2 and 4-1BB surface protein expression, in which the cells were stained with the same near-infrared live/dead stain (#L34975, Invitrogen) in PBS, followed by resuspension in PBS containing 2% FBS for staining with ST2-BV605 (#745257, BD Biosciences) and 4-1BB-AlexaFluor647 (#FAB937R, R&D Systems) for 30 minutes on ice, washed and resuspended in PBS containing 2% FBS (#SH30071.03, Hyclone, GE Life Sciences) prior to analyzing in triplicates.

#### RNA extraction and bulk RNA-seq library preparation

Trizol (#15596018, Invitrogen) was added to each Treg cell sample containing approximately 50,000 murine splenic Treg cells, which were stored at −80°C until RNA extraction. RNA extraction was performed according to the manufacturer’s protocol. In brief, chloroform was added, and the sample was centrifuged, giving a lower red phenol-chloroform layer, an interphase, and a colorless upper aqueous phase. The colorless aqueous phase containing RNA was collected. The RNA was precipitated with isopropanol and washed with 75% ethanol. After subsequent drying, the pellet was resuspended in RNase-free water. RNA quality was assessed by TapeStation 4200 (Agilent Technologies) and Nanodrop 8000 spectrophotometer (Thermo Scientific), and RNA quantity was determined by Qubit 4 fluorometer (Invitrogen). Bulk RNA-seq libraries were prepared according to the Illumina TruSeq stranded mRNA (#20020595, Illumina) workflow with modifications. ERCC spiked-in 150 µg of RNA was used as input, which was subjected to purification by binding to RNA binding beads. The purified RNA was fragmented, followed by first and second strand synthesis, A-tailing, adapter ligation and PCR library amplification. Final RNA libraries were subjected to qualitative and quantitative quality control by the TapeStation 4200 and Qubit 4, respectively, prior to pooling and sequencing on a NextSeq2000 at PE100.

#### Bulk ATAC-seq library preparation

Bulk ATAC-seq libraries, each with approximately 50,000-100,000 Treg cells, were prepared using the Active Motif ATAC-seq kit (#105320, Active Motif). Nuclei were extracted from the freshly isolated and sorted double positive and double negative ST2 and 4-1BB protein expressing Treg cells using the provided ATAC lysis buffer, open chromatin regions were tagmented with adapters, tagmented DNA was purified and amplified, and libraries were cleaned-up with SPRI beads. Final libraries were assessed qualitatively by TapeStation 4200 (Agilent Technologies) and quantitatively by Qubit 4 fluorometer (Invitrogen) prior to pooling and sequencing on a NextSeq2000 at PE50.

#### Bulk RNA-seq data analysis

Raw RNA sequencing FASTQ reads were aligned to the mouse reference (B39) using Qiagen’s Omicsoft implemented OSA aligner. Raw gene level counts were determined using Omicsoft’s RSEM implementation and relying on the GENCODE v28 gene model. Downstream analysis was performed using R (version 4.4.0). Differential gene expression analysis between ST2^+^4-1BB^+^ (activated) and ST2^-^4-1BB^-^ (resting) cells was performed using DESeq2 (version 1.44.0) (Love et al., 2014). P-values were adjusted by apeglm shrinkage. The significant (adjusted p-value < 0.05) DEGs were used as input into Ingenuity Pathway Analysis (IPA) to generate the pathway enrichment bar graph.

#### Bulk ATAC-seq data analysis

ATAC-Seq was processed using nf-core/atacseq pipeline (10.5281/zenodo.2634132) to get consensus peaks for downstream analysis. Mouse reference genome cellranger-arc-mm10-2020-A-2.0.0 provided by 10x Genomics was used as reference for read mapping. Differentially accessible peaks between activated and resting Treg cells were obtained using DESeq2 (version 1.44.0). P-values were adjusted by apeglm shrinkage. Peaks were annotated using ChIPseeker (version v1.40.0) (Wang et al., 2022a, Yu et al., 2015) with defined TSS region as range from −1000bp to 100bp and genes were annotated with TxDb.Mmusculus.UCSC.mm10.knownGene (Team and Maintainer, 2019). Motifs from the JASPAR database (Baranasic, 2022) that associated with variability in chromatin accessibility between activated and resting Treg samples were identified using ChromVar (Schep et al., 2017).

##### Differential footprinting for ATAC-seq

In order to identify the differential TF activity between Treg cells in resting and activated states, we compared the footprints of the activated and resting Tregs using HINT-ATAC (Li et al., 2019). Differential analysis was performed to obtain TF activity score, tag counts and p-value for each TF.

##### Classification of transcriptional activators and repressors

In order to quantify the difference in TF activity and to classify TFs as putative transcriptional activators or repressors, we applied diffTF (Berest et al., 2019) to the RNA-seq and ATAC-seq data using PWM database PWMScan_HOCOMOCOv10 (Ambrosini et al., 2018).

#### Nuclei isolation and 10x multiome

Immunomagnetically enriched murine Treg cells were extracted for nuclei according to the low input 10x nuclei extraction protocol for the Multiome assay (#CG000365 Rev C, 10x Genomics). In brief, cell membranes were lysed in a lysis buffer containing detergents, such as 0.1% Tween-20 (#1662404, Bio-Rad), 0.1% Nonidet P40 (NP40) (#74385, Sigma-Aldrich) and 0.01% digitonin (#BN2006, Thermo Fisher), for 30 seconds on ice, followed by neutralization with a wash buffer that consists of 1% BSA. The final nuclei pellet was resuspended in the 10x Genomics provided diluted nuclei buffer with 1 unit/uL RNase inhibitor (#3335402001, Sigma-Aldrich) and 1 mM dithiothreitol (DTT). Manual nuclei count in trypan blue on a hematocytometer determined the targeted sequencing nuclei recovery. This was followed by transposition, GEM generation and barcoding, and pre-amplification, according to the manufacturer’s protocol for 10x Genomics Next GEM Single Cell Multiome ATAC and Gene Expression assay (#CG000338 Rev F, 10x Genomics). Each pre-amplified sample was split for downstream ATAC library construction or cDNA amplification for 3’ gene expression library construction. The snATAC-seq libraries were prepared manually according to the manufacturer’s protocol (#CG000338 Rev F, 10x Genomics), while for the snRNA-seq libraries, the cDNA was amplified manually according to the protocol (#CG000338 Rev F, 10x Genomics) and the remaining gene expression libraries were constructed robotically using the Chromium Connect (#CG000286 Rev E, 10x Genomics). All libraries were assessed qualitatively by TapeStation 4200 (Agilent Technologies) and quantitatively by Qubit 4 fluorometer (Invitrogen) prior to pooling the snATAC-seq and snRNA-seq libraries separately and sequencing on a NovaSeq6000.

#### 10x Multiome data analysis

##### Pre-processing of the 10x multiome snRNA/ATAC-seq

Raw FASTQ sequencing reads from both gene expression and DNA accessibility libraries were processed using 10x Genomics Cell Ranger ARC v2.0.0. The mouse reference genome, cellranger- arc-mm10-2020-A-2.0.0, provided by 10x Genomics was used as reference for read mapping. UMI counts (GEX) and fragments (ATAC) were outputs for downstream analysis using Seurat (version 5.1.0) (Hao et al., 2024) and Signac (1.13.0) (Stuart et al., 2021). In order to remove low quality cells, cells were kept based on nCount_ATAC (>1000 and < 80000) and nucleosome_signal (<2) and TSS.enrichment (>2).

##### Peak, gene activity and chromVAR from snATAC-seq

Peak assay was derived from peak calling on snATAC-Seq using MACS2 (Zhang et al., 2008a). Peaks were annotated using ChIPseeker with defined TSS region as range from −1000bp to 100bp and genes were annotated with as TxDb.Mmusculus.UCSC.mm10.knownGene.

Since we have low snRNA-Seq coverage, we generated gene activity assay from snATAC-seq as an alternative to gene expression assay from snRNA-seq to enable downstream gene expression analysis.

In order to quantify motif activity in each cell and compare motif activities between cell states, we generated a motif activity score assay using the RunChromVAR function in Seurat based on chromVAR (Schep et al., 2017).

##### Normalization, dimensionality reduction, clustering and visualization

Normalization, dimensionality reduction, cell clustering and visualization were performed using Seurat and Signac.

For the RNA assay, the count matrix was normalized using log-transformation and scaled using Seurat. The top 3000 variable features were selected. Principle component analysis (PCA) was performed using 50 principal components. The first 30 principal components were used to find k- nearest neighbors (k=20). Unsupervised clusters were derived using the original Louvain algorithm with resolution=0.3. Cell clustering was visualized using UMAP (uwot method) (McInnes et al., 2018).

For the peak assay, latent semantic indexing (LSI), which included a combination of frequency- inverse document frequency (TF-IDF) and singular value decomposition (SVD), was performed. First, the count matrix was normalized using TF-IDF method. Next, SVD was performed on the TD- IDF matrix to obtain a reduced dimension representation. The 2nd to 30th dimensions from LSI reduction were used to find k-nearest neighbors (k=20), since the first dimension is typically correlated with sequencing depth. Unsupervised clusters were derived using SLM algorithm with resolution=0.3. Cell clustering was visualized using UMAP (uwot method).

For the gene activity assay, the count matrix was normalized using log-transformation and scaled using Seurat. The top 3000 variable features were selected. Principle component analysis (PCA) was performed using 50 principal components. The first 30 principal components were used to find k-nearest neighbors (k=20). Unsupervised clusters were derived using the original Louvain algorithm with resolution=0.3. Cell clustering was visualized using UMAP (uwot method).

##### Cell type prediction

To infer the cell states, we performed label transfer from a reference scRNA-seq dataset (Lu et al., 2020) to the gene activity assay for spleen and lung separately using Seurat. The reference scRNA- seq was first log-transformed and scaled. Next, a set of anchors between the reference scRNA-seq and the gene activity assay using the FindTransferAnchors function with cca reduction was determined. Then, the cell state labels from the reference dataset were transferred to the cells in our study using the TransferData function (weight reduction=LSI and dimensions=2:30).

##### Linking peaks to genes

In order to find peaks that were correlated to the expression of nearby genes, we linked peaks to gene activity of nearby genes using the LinkPeaks function in Seurat.

##### Transcription factor footprinting

To compare the TF footprints in different cell states, we first added the JASPAR2020 motif matrix to the peak assay and ran the Footprint function in Seurat to retrieve the footprints of all cell states for each TF. Next, the cells were down-sampled to allow a maximum of 1000 cells per cell state.

##### Differential analysis

Differential gene expression analysis was performed using the Seurat functions FindAllMarkers (for multiple groups) and FindMarkers (for two groups) using the gene activity assay. Differential motif activity analysis was also performed using the Seurat functions FindAllMarkers (for multiple groups) and FindMarkers (for two groups) using the motif activity assay. Differential gene expression and motif activity were reported based on fold change and adjusted p-value.

#### DBiT-seq based spatial ATAC-seq assay and data analysis

FOXP3-GFP^+^ mice were injected intraperitoneally with PBS or mIL-2M for four days, after which the lower left lungs were isolated and subjected to simultaneous freezing and embedding with OCT (#4583, Sakura) in isopentane (#270342, Sigma-Aldrich) and liquid nitrogen. Tissues were stored in −80°C until sectioned. Frozen lung blocks were cryosectioned at 10 µm thickness and transferred onto Superfrost Plus microscope glass slides (#1255015, Fisher Scientific) for DBiT-seq based spatial ATAC-seq assay using the 3.8 mm x 3.8 mm chip comprising of 9216 tixels with 20 µm resolution. Targeted sequencing per sample was 1.1 billion PE reads.

Fragment files were processed and analyzed using the ArchR pipeline (Granja et al., 2021). First, Arrow files were created for each sample and aligned to the mm10 genome to generate ArchR projects. Then, spatial coordinates and off-tissue tixels were filtered using AtlasXomics’ addition to the original ArchR pipeline, followed by LSI dimensionality reduction using 25,000 variable features and clustering at low resolution to annotate cell niches based on significant (FDR < 0.05) cluster markers with high differential gene scores relative to all other clusters. Seurat objects were built from the ArchR projects for spatial cluster and gene visualizations.

#### Source and in vitro activation of human Treg cells

Freshly isolated healthy adult human Treg cells from peripheral blood mononuclear cells (PBMCs) were purchased from Human Cells Biosciences (California, USA). Naïve/unstimulated cells were used immediately for the CUT&RUN and bulk ATAC-seq assays. For activated/stimulated cells, freshly isolated cells were allowed to recover in X-vivo 20 culture medium (#04-448Q, Lonza) containing 5% heat-inactivated fetal bovine serum (FBS) (#SH30071.03, Hyclone, GE Life Sciences) and 100 units/mL of human recombinant IL-2 (#589102, BioLegend) for two days at 37°C with 5% CO_2_, followed by stimulation with human anti-CD3 and anti-CD28 activation beads (#10971, StemCell Technologies) for two days and 500 units/mL of human recombinant IL-2 (#589102, BioLegend) for an additional three days prior to proceeding with the CUT&RUN and bulk ATAC-seq assays.

#### Human Treg CUT&RUN and bulk ATAC-seq

##### CUT&RUN assay, data analysis and visualization

Freshly isolated naïve and recombinant human IL-2 stimulated human PBMC Treg cells (#PBCD4-Treg, Human Cells Biosciences) were subjected to nuclei isolation, followed by the CUT&RUN kit (#SKU14-1048, Epicypher) was used to prepare sequence-ready libraries. In brief, the nuclei were immobilized onto beads and permeabilized to allow for antibody binding. Then, a fusion of Protein A to Micrococcal Nuclease (pA-MNase) was added to cleave antibody-bound chromatin. Testing antibodies utilized included anti-BACH1 (#66762-1-Ig, Proteintech and # A303-057A, Bethyl Laboratories, anti-BATF (#pab4003, Brookwood Medical and # ab236876, Abcam), and controls including anti-H3K4me3 (#13-0041, Epicypher), anti-H3K27ac (#13-0045, Epicypher), anti-H3K9me3 (#ab176916, Epicypher) and anti-IgG (#13-0042, Epicypher). Adapters were added to the chromatin DNA fragments, and Illumina short read sequencing was performed.

CUT&RUN data were processed using the nf-core (Ewels et al., 2020) CUT&RUN pipeline, version 3.2.1 (Cheshire et al., 2023). For the most part, we used default parameters, but CPM, rather than spike-in, was chosen for normalization, given relatively variable spike-in alignment rates across samples. Reads were aligned to hg38 using a macs_gsize of 2.8 E9, and peaks were identified for each assayed factor (BACH1 and BATF) using seacr (Meers et al., 2019).

Identified peaks were further interrogated with HOMER, version 4.11.1 (Heinz et al., 2010), to find transcription factor (TF) motifs enriched at the binding sites of our assayed factors. Specifically, we used the findMotifsGenome.pl script on bed-formatted files of identified peaks, setting hg38 as the reference genome. The top 20 most significant known TF motifs enriched in each condition, and their parent categories, were then assessed and compared. We further interrogated the identified peaks using the “peakHeatmap”, “peak_Profile_Heatmap”, and “plotAnnoPie” functions from ChIPseeker, version 1.40.0 (Wang et al., 2022b), to ascertain genomic annotations at identified peaks, as well as to visualize binding signal profile across gene bodies and transcription start sites (TSSs). Peak annotations were additionally assessed using the annotatr package, version 1.30.0 (Cavalcante and Sartor, 2017), to include additional finer-grained gene and enhancer features on the hg38 genome.

##### Bulk ATAC-seq, data analysis and visualization

ATAC-seq data were processed using the nf-core (Ewels et al., 2020) ATAC-seq pipeline, version 2.1.2 (Patel et al., 2023). The pipeline’s default parameters were used wherever applicable, and we aligned reads to GRCm39 using a macs_gsize of 1.87 E9 and narrow peak calling with MACS2 (Zhang et al., 2008b). Read signal and identified peaks in each condition were visualized using integrative genomics viewer (Robinson et al., 2011).

#### Gene editing by CRISPR knockout

##### Ex vivo CRISPR knockout

FACS sorted mIL-2M stimulated mouse splenic FOXP3-GFP^+^ Tregs (with about 500,000 cells per reaction) were resuspended in supplemented P4 solution (#PBP4-00675, Lonza Bioscience) and incubated for 15 minutes with ribonucleoprotein (RNP) complexes, comprising of 60 pmol Cas9-RFP (#10008163, IDT) and a single or dual pool of 100 pmol of each of three single-guide ribonucleic acids (sgRNAs) (Synthego) per gene, followed by addition of an electroporation enhancer (#10007805, IDT) at 4 µM final concentration and electroporation using the Lonza Bioscience 4D Nucleofector system and the DS-137 program in a strip format.

The stimulated control consisted of Cas9-RFP only without sgRNAs and was electroporated. The cell-RNP complexes were in a 25 µL solution. After an incubation of 3 minutes, each cell-RNP solution was transferred to a well of a 16-well strip (#V4XP-4032, Lonza) and immediately electroporated. Cells were allowed to recover for 10 minutes post-electroporation at room temperature, at the end of which 80 µL of complete medium of Advanced RPMI1640 (#12633-012, Gibco) containing 10% heat-inactivated FBS (#SH30071.03, Hyclone, GE Life Sciences), 0.1% β-mercaptoethanol (#21985-023, Gibco), 2 mM L-Glutamine, (#25030-081, Gibco), 100 units/mL penicillin and 100 µg/mL streptomycin (#15140-122, Gibco) and 100 units/mL of human recombinant IL-2 (#589102, BioLegend) was added. Thereafter, 100 µL of the cell solution was transferred to each of a well of a 6-well plate containing the same complete medium prewarmed to 37°C. The cells were stimulated with anti-CD3 and anti-CD28 activation beads (#11452D, Invitrogen) at a bead to cell ratio of one-to-one and allowed to be stimulated and cultured for 3 days for gene editing to occur. At the end of the 3 days, cells were replenished with fresh medium, replacing human recombinant IL-2 with 1 nM mIL-2M for stimulation, and cultured for another 3 days, at the end of which flow cytometry, bulk ATAC-seq and genomic DNA extraction for ICE analysis were performed. The method details for flow cytometry and bulk ATAC-seq were outlined above.

#### CRISPR Genomic DNA extraction and editing efficiency

The DNeasy Blood and Tissue kit using spin columns (#69504, Qiagen) was used to isolate genomic DNA (gDNA) from the Treg cells. In brief, supernatant removed pelleted cells upon centrifugation were lysed in a lysis buffer in the presence of RNase A and proteinase K. Lysed cell debris was removed by rendering genomic DNA a negative charge upon mixing with absolute ethanol to bind to the column membrane while the cell debris filtered through. The spin columns were washed and the pure gDNA was eluted. The gDNA was used as a template for amplification of CRISPR target genes using PCR primers. Gel electrophoresis using TapeStation 4200 (Agilent Technologies) D5000 was used to assess for the presence of indels in the amplified DNA. Sanger sequencing was performed on the purified amplicons using the listed sequencing primers. Interference of CRISPR Edits (ICE) analysis was carried out on the .abl files generated from Sanger sequencing to calculate the CRISPR knockout efficiencies for the tested gene edits.

### Quantification and statistical analysis

For the statistical analysis used to quantify data, please refer to the STAR Methods sub-sections above. Statistical parameters are reported in the STAR Methods and Figure legends. One-way ANOVA followed by Dunnett’s multiple comparison test were performed using GraphPad Prism (v10.2.3). All computational analyses were performed using R version 4.4.0.

## Supplemental Information Titles and Legends

Document S1. Figures S1-S14.

Table S1. Bulk RNA-seq z-scored normalized counts, related to Figure 1.

Table S2. Bulk RNA-seq differentially expressed genes list, related to Figure 1C.

Table S3. Bulk ATAC-seq annotated differentially accessible peaks, related to Figure 1.

Table S4. Bulk ATAC-seq enriched motifs, related to Figure 1G.

Table S5. Single-nuclei ATAC-seq annotated chromatin accessibility based on samples, related to Figure 2.

Table S6. Single-nuclei ATAC-seq annotated chromatin accessibility based on cell state, related to Figure 2.

Table S7. Single-nuclei ATAC-seq transcription factor motifs enrichment based on cell state, related to Figure 2.

Table S8. Single-nuclei ATAC-seq significant transcription factor motifs enrichment based on cell state, related to Figure 2.

Table S9. Single-nuclei ATAC-seq differential transcription factor motifs between cell states, related to Figure 2.

Table S10. Single-nuclei RNA-seq downstream genes of motifs overlapping between clusters/cell states, related to Figure 2.

Table S11. Lung single-nuclei ATAC-seq annotated chromatin accessibility based on samples, related to Figure 3C.

Table S12. Lung single-nuclei ATAC-seq annotated chromatin accessibility based on cell states, related to Figure 3D.

Table S13. Lung spatial ATAC-seq cluster differential accessible gene score, related to Figure 4E.

Table S14. Differential ATAC-seq peaks for selective bZIP perturbation, related to Figure 5.

Table S15. Human bulk ATAC-seq enriched transcription factor motifs, related to Figure 6.

Table S16. BATF and BACH1 binding targets by CUT&RUN in human naïve and activated Treg, related to Figure 6.

Table S17. GO analysis of nearest gene downstream of promoter annotated peaks from BATF and BACH1 CUT&RUN, related to Figure 6.

