## Supplementary Figures for "Basic-Leucine-Zipper Transcription Factors Regulate Selective Molecular Phenotypes in Regulatory T Cells During IL-2-Induced Activation"

(Supplementary) Figure S1

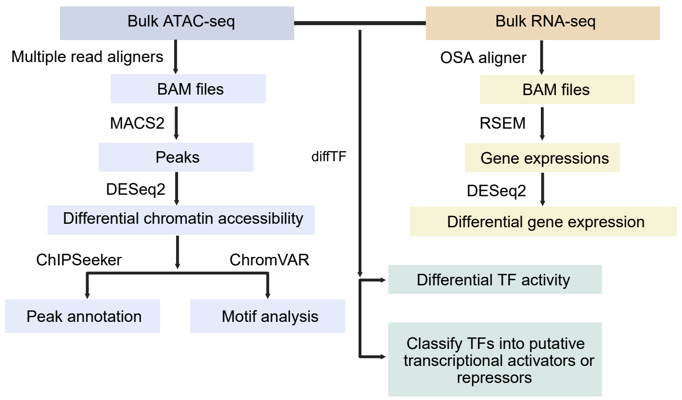

(Supplementary) Figure S2

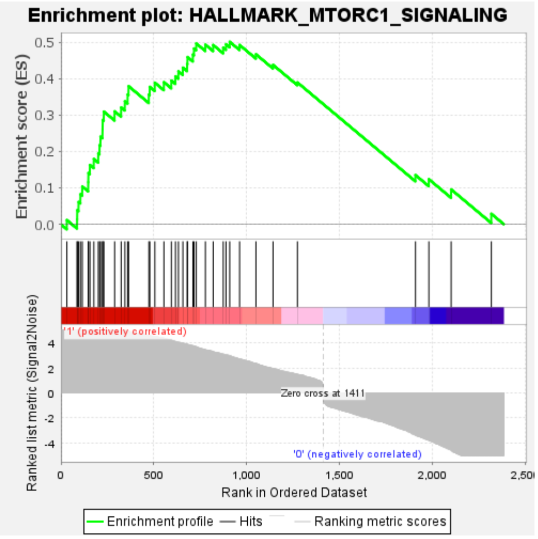


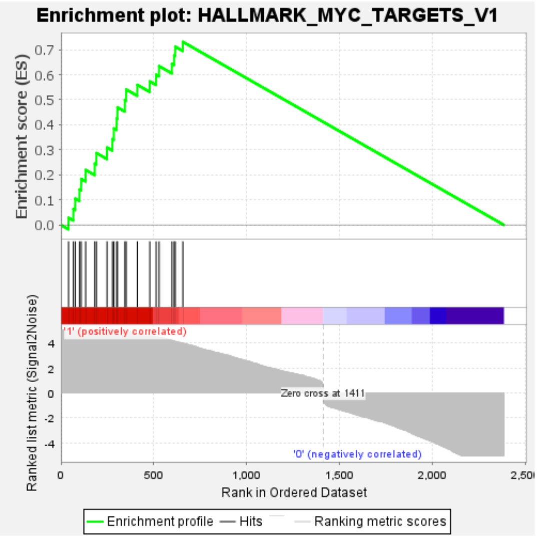


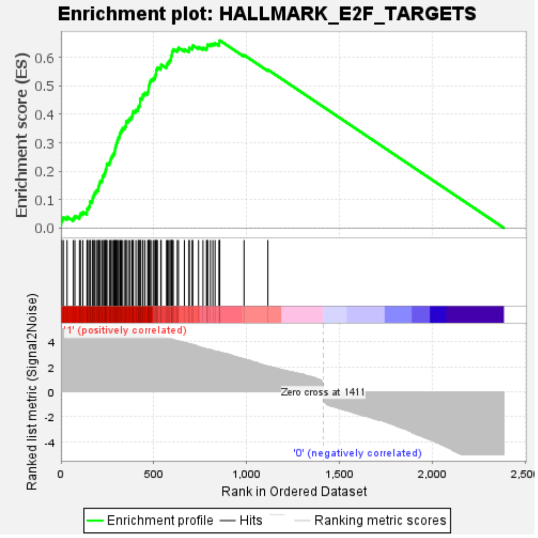


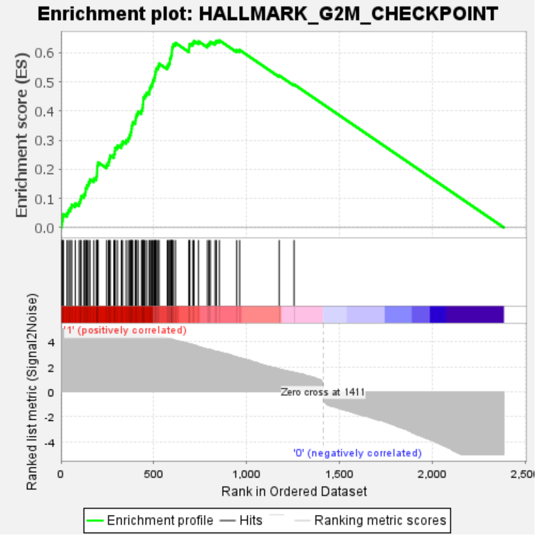


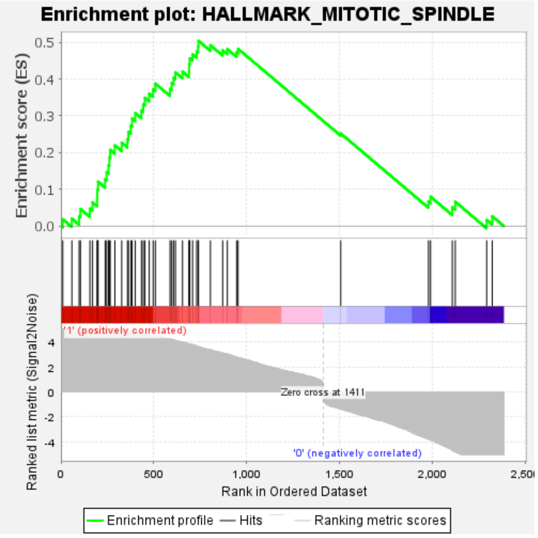

(Supplementary) Figure S3

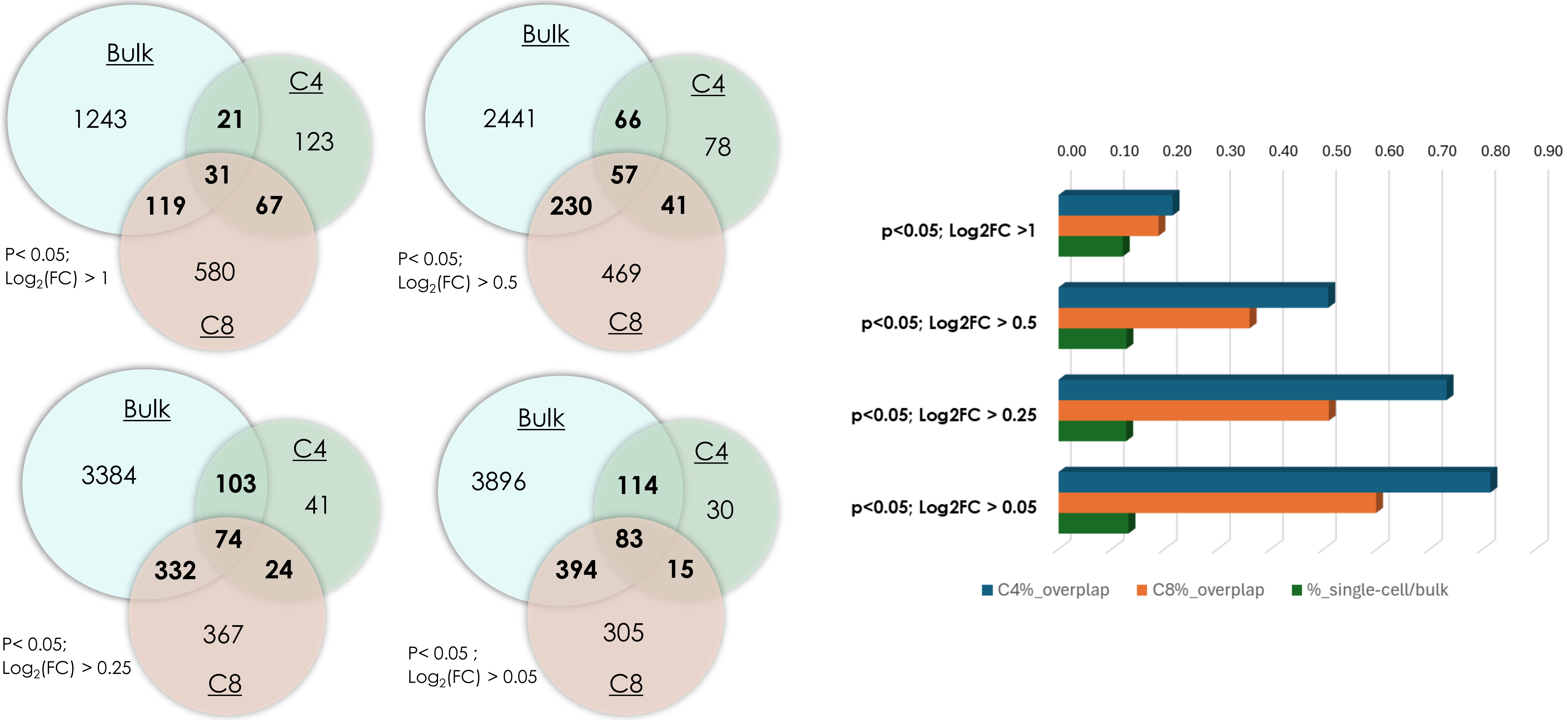



Internal Use Only General and Administrative

AMGEN

(Supplementary) Figure S4

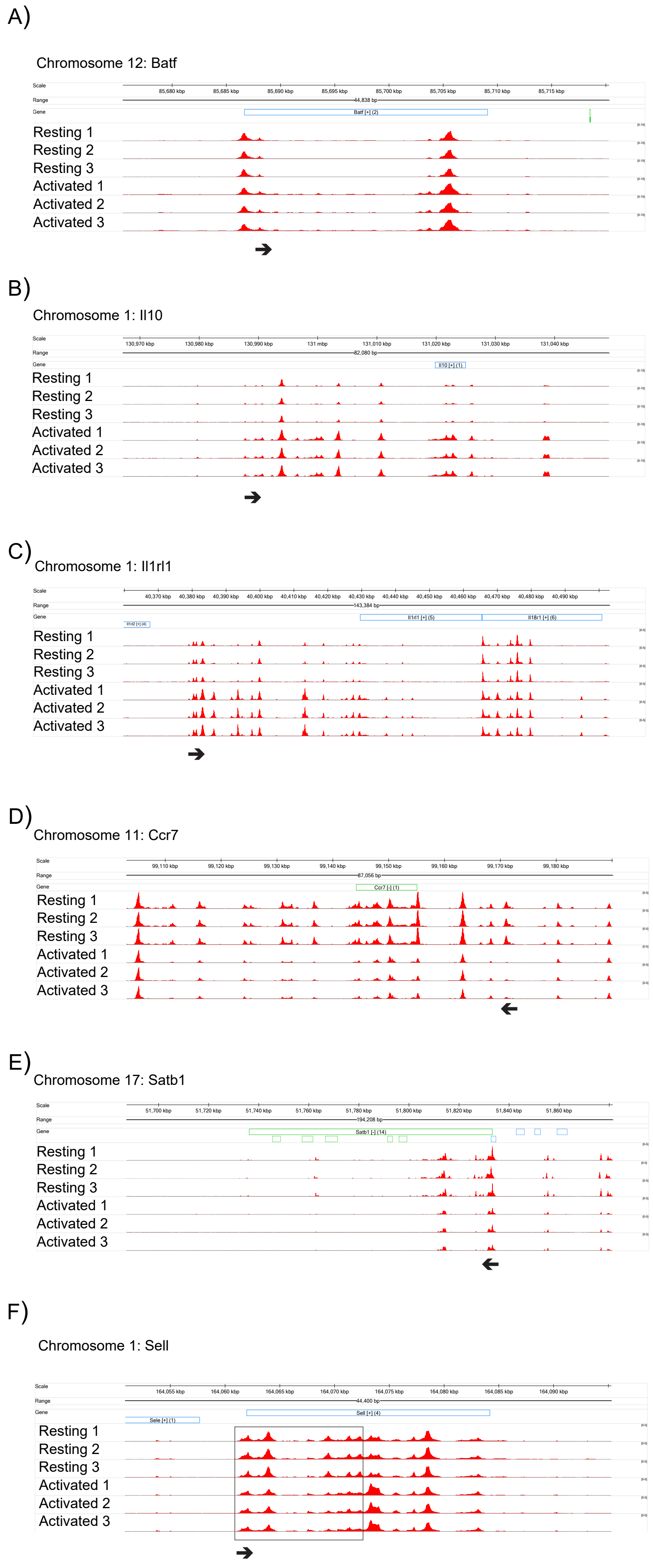

(Supplementary) Figure S5

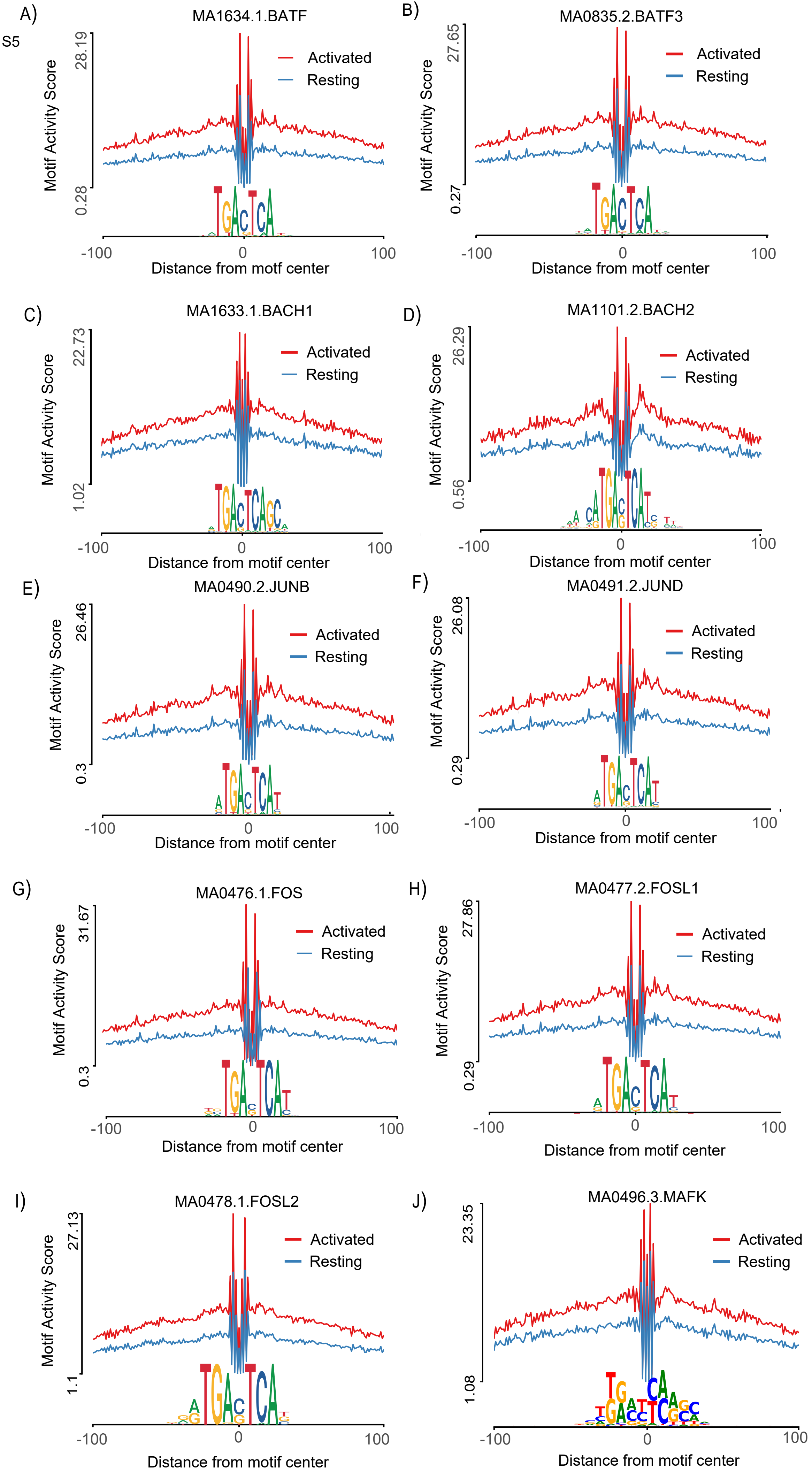

(Supplementary) Figure S6

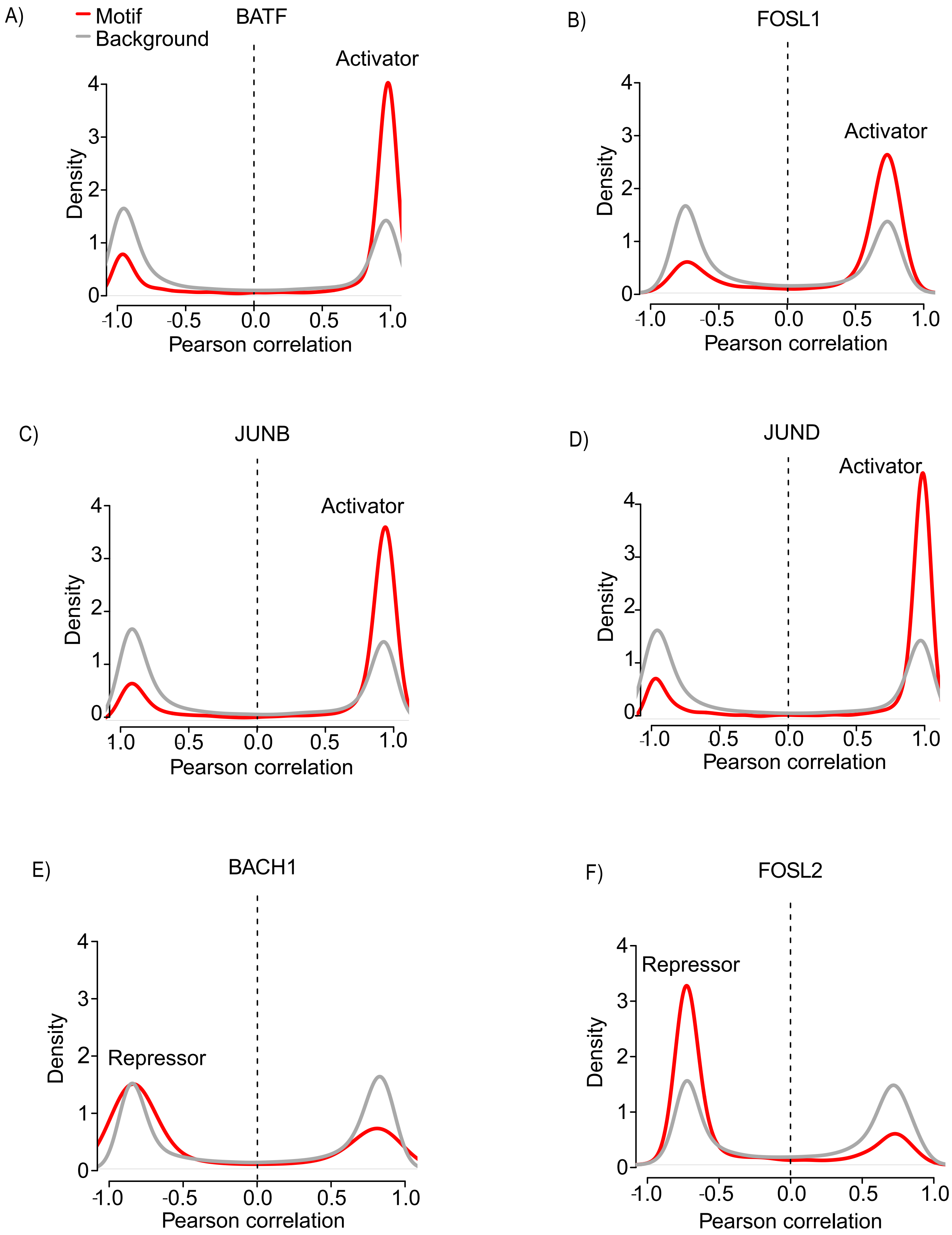

(Supplementary) Figure S7

A)

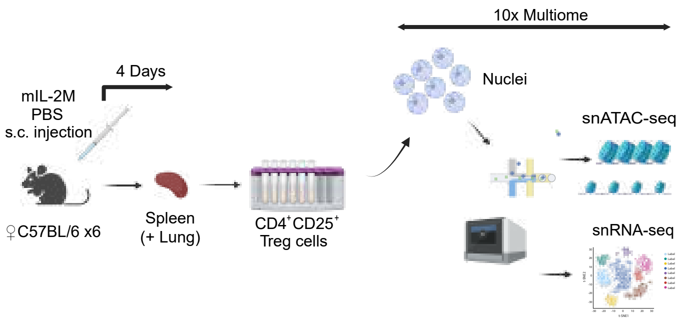


B)

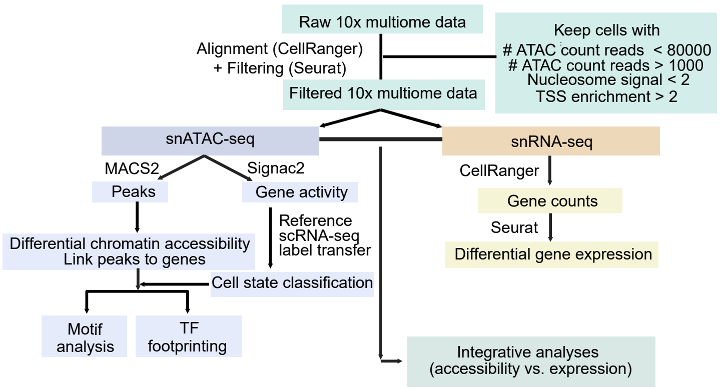

(Supplementary) Figure S8

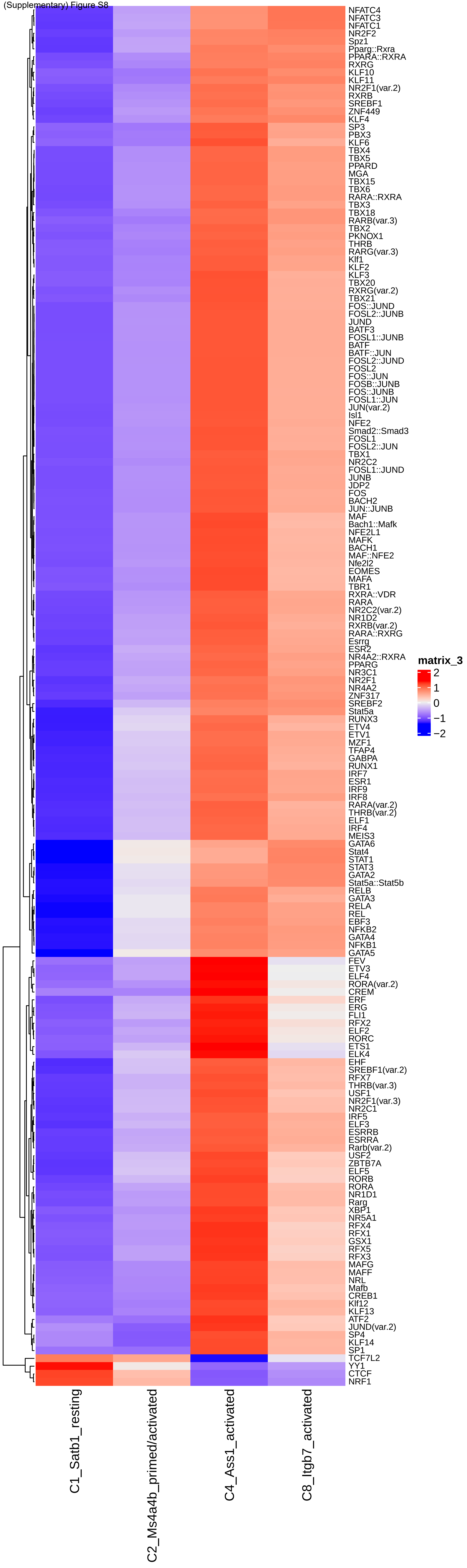

(Supplementary) Figure S9

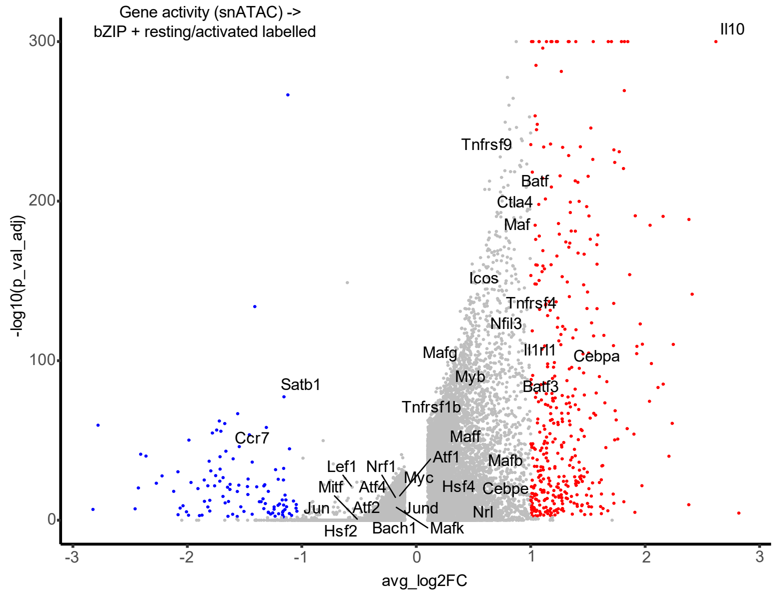


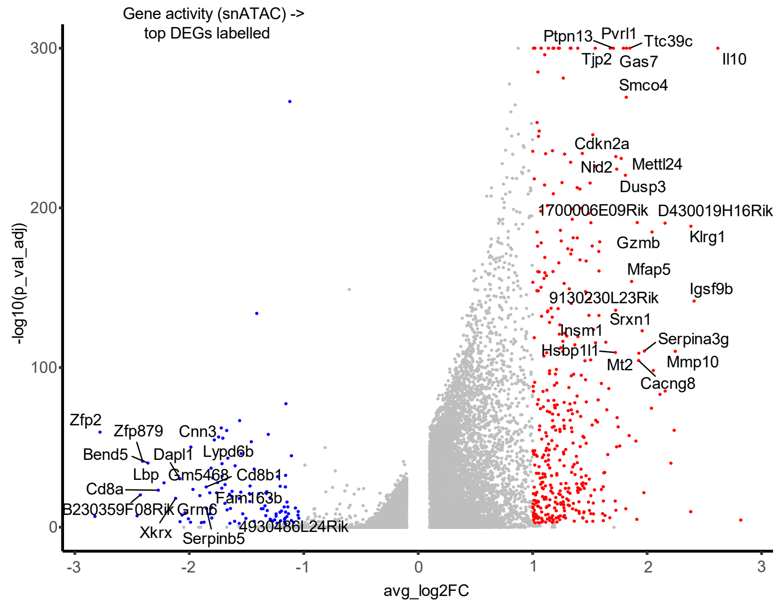


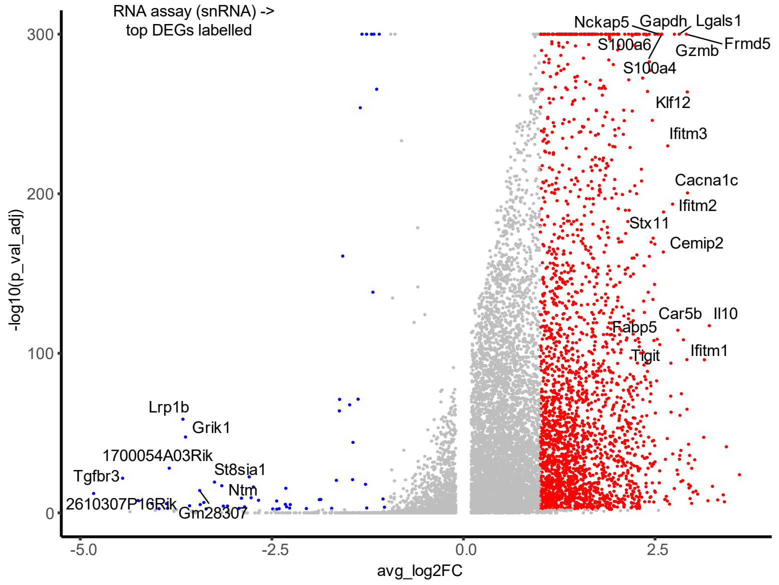

(Supplementary) Figure S10

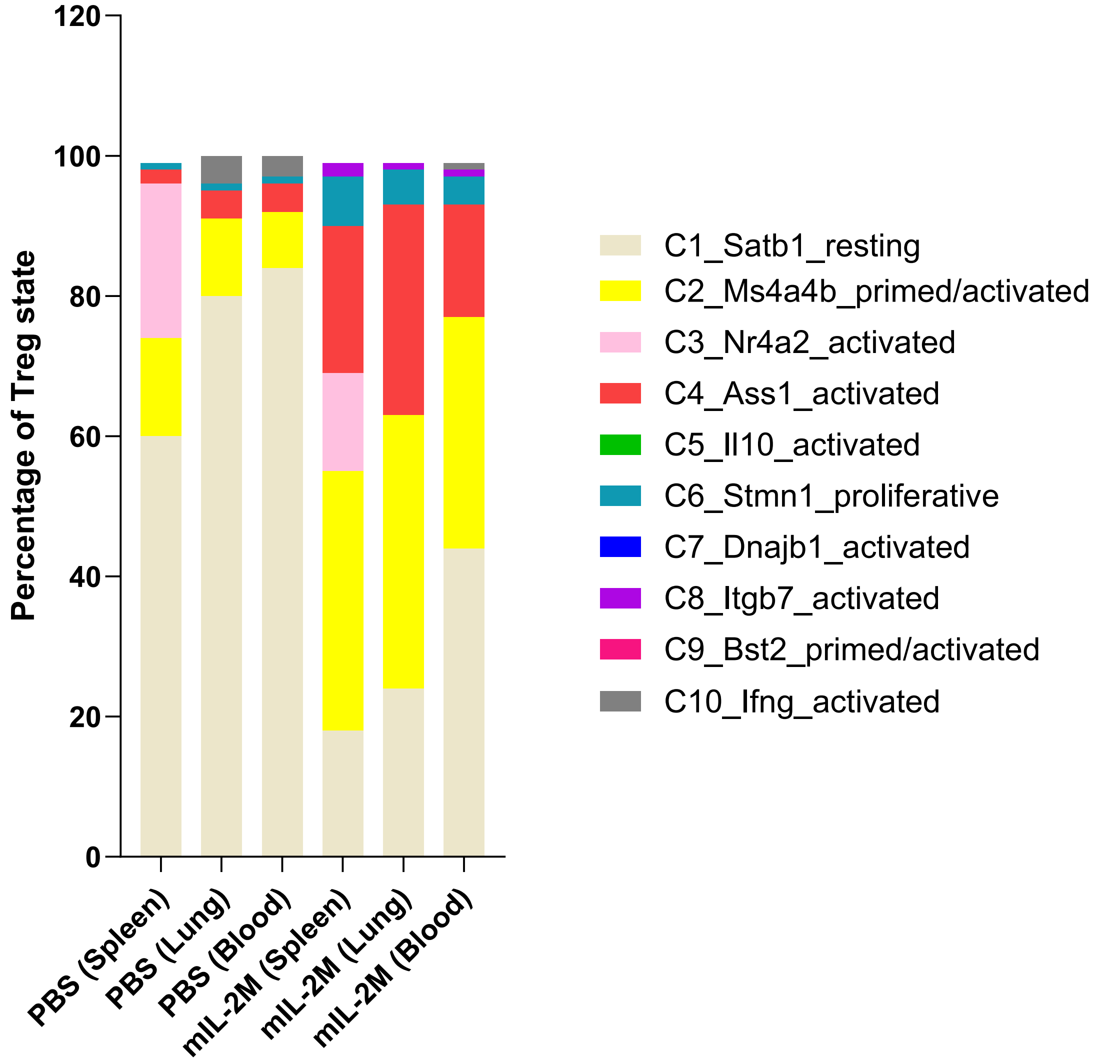

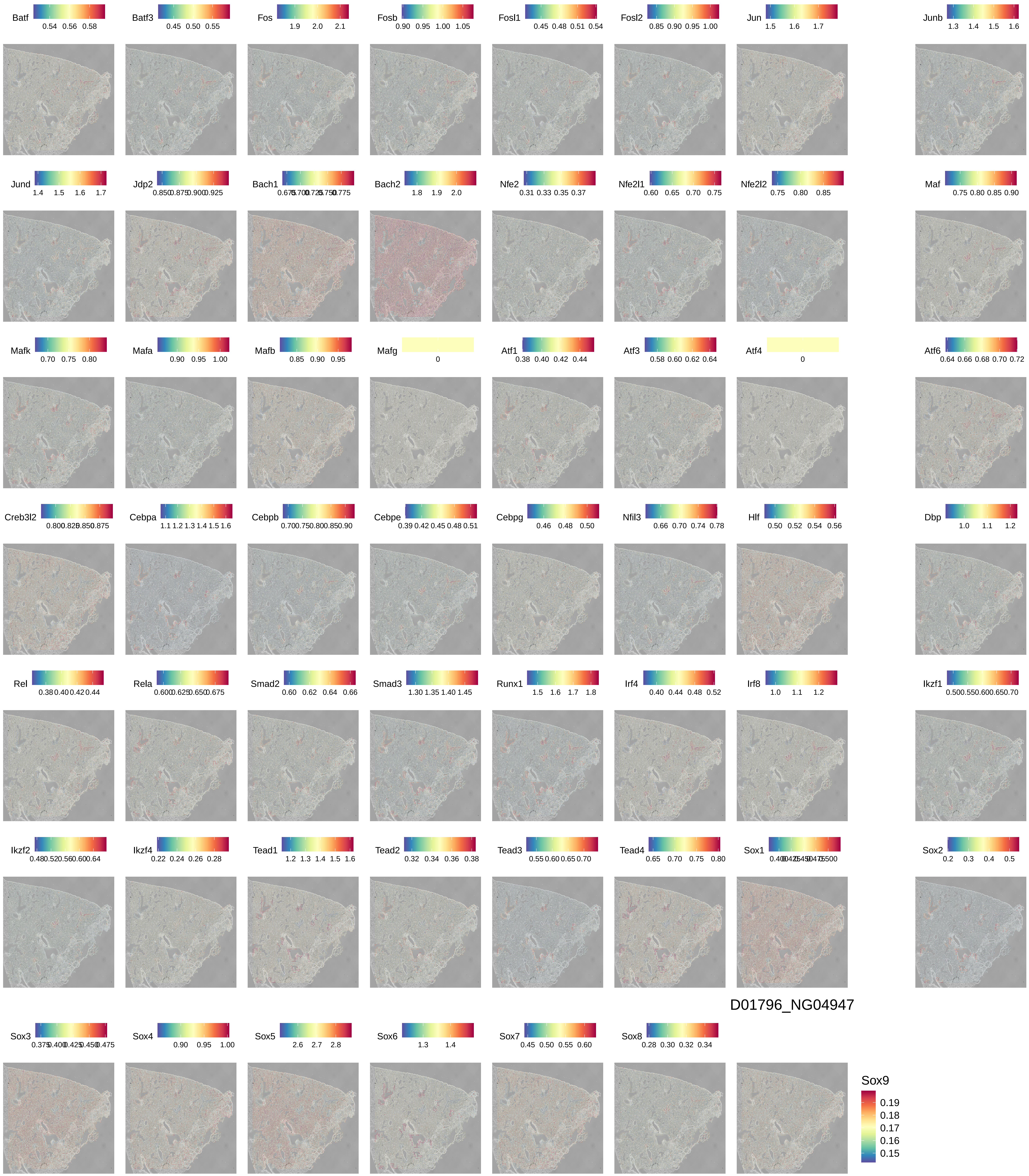

Batf 0.54 0.56 0.58
Batf3 0.45 0.50 0.55
Fos 1.9 2.0 2.1
Fosb 0.90 0.95 1.00 1.05
Fosl1 0.45 0.48 0.51 0.54
Fosl2 0.85 0.90 0.95 1.00
Jun 1.5 1.6 1.7
Junb 1.3 1.4 1.5 1.6
Jund 1.4 1.5 1.6 1.7
Jdp2 0.850 0.875 0.900 0.925
Bach1
Bach2 1.8 1.9 2.0
Nfe2 0.31 0.33 0.35 0.37
Nfe2l1 0.60 0.65 0.70 0.75
Nfe2l2 0.75 0.80 0.85
Maf 0.75 0.80 0.85 0.90
Mafk 0.70 0.75 0.80
Mafa 0.90 0.95 1.00
Mafb 0.85 0.90 0.95
Mafg 0
Atf1 0.38 0.40 0.42 0.44
Atf3 0.58 0.60 0.62 0.64
Atf4 0
Atf6 0.64 0.66 0.68 0.70 0.72
Creb3l2
Cebpa 1.1 1.2 1.3 1.4 1.5 1.6
Cebpb
Cebpe 0.39 0.42 0.45 0.48 0.51
Cebpg 0.46 0.48 0.50
Nfil3 0.66 0.70 0.74 0.78
Hlf 0.50 0.52 0.54 0.56
Dbp 1.0 1.1 1.2
Rel 0.38 0.40 0.42 0.44
Rela 0.600 0.625 0.650 0.675
Smad2 0.60 0.62 0.64 0.66
Smad3 1.30 1.35 1.40 1.45
Runx1 1.5 1.6 1.7 1.8
Irf4 0.40 0.44 0.48 0.52
Irf8 1.0 1.1 1.2
Ikzf1
Ikzf2
Ikzf4 0.22 0.24 0.26 0.28
Tead1 1.2 1.3 1.4 1.5 1.6
Tead2 0.32 0.34 0.36 0.38
Tead3 0.55 0.60 0.65 0.70
Tead4 0.65 0.70 0.75 0.80
Sox1
Sox2 0.2 0.3 0.4 0.5
D01796_NG04947
Sox3
Sox4 0.90 0.95 1.00
Sox5 2.6 2.7 2.8
Sox6 1.3 1.4
Sox7 0.45 0.50 0.55 0.60
Sox8 0.28 0.30 0.32 0.34
Sox9 0.19 0.18 0.17 0.16 0.15

(Supplementary) Figure S12

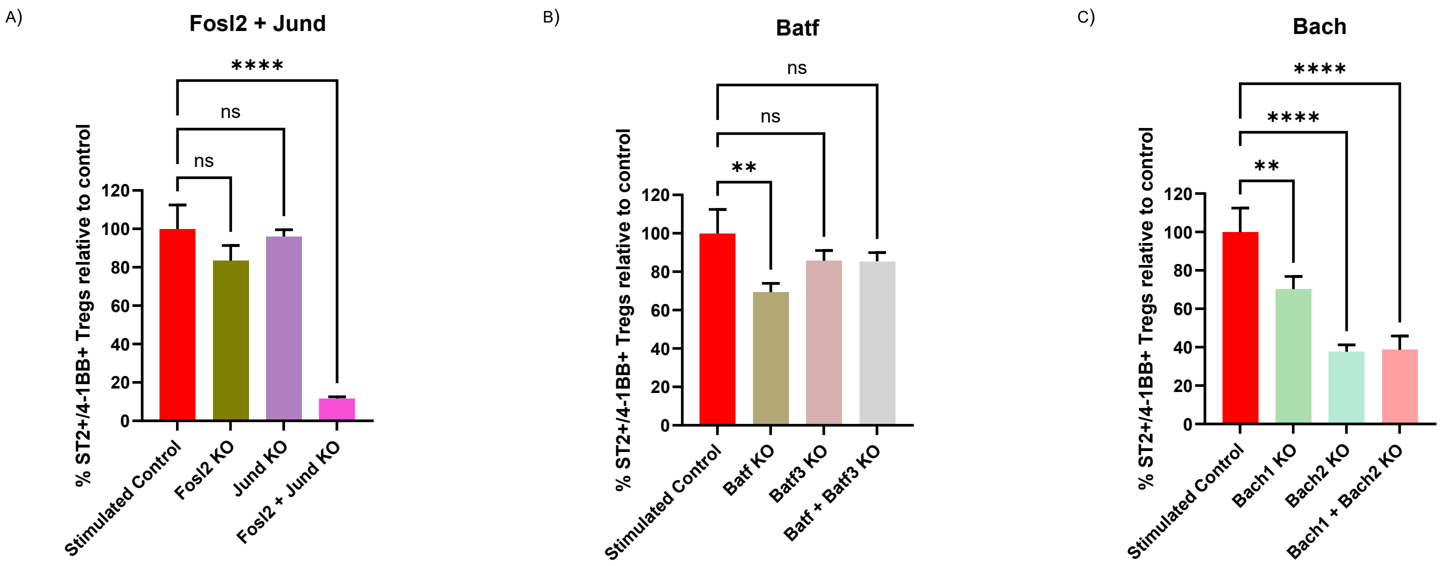

(Supplementary) Figure S13

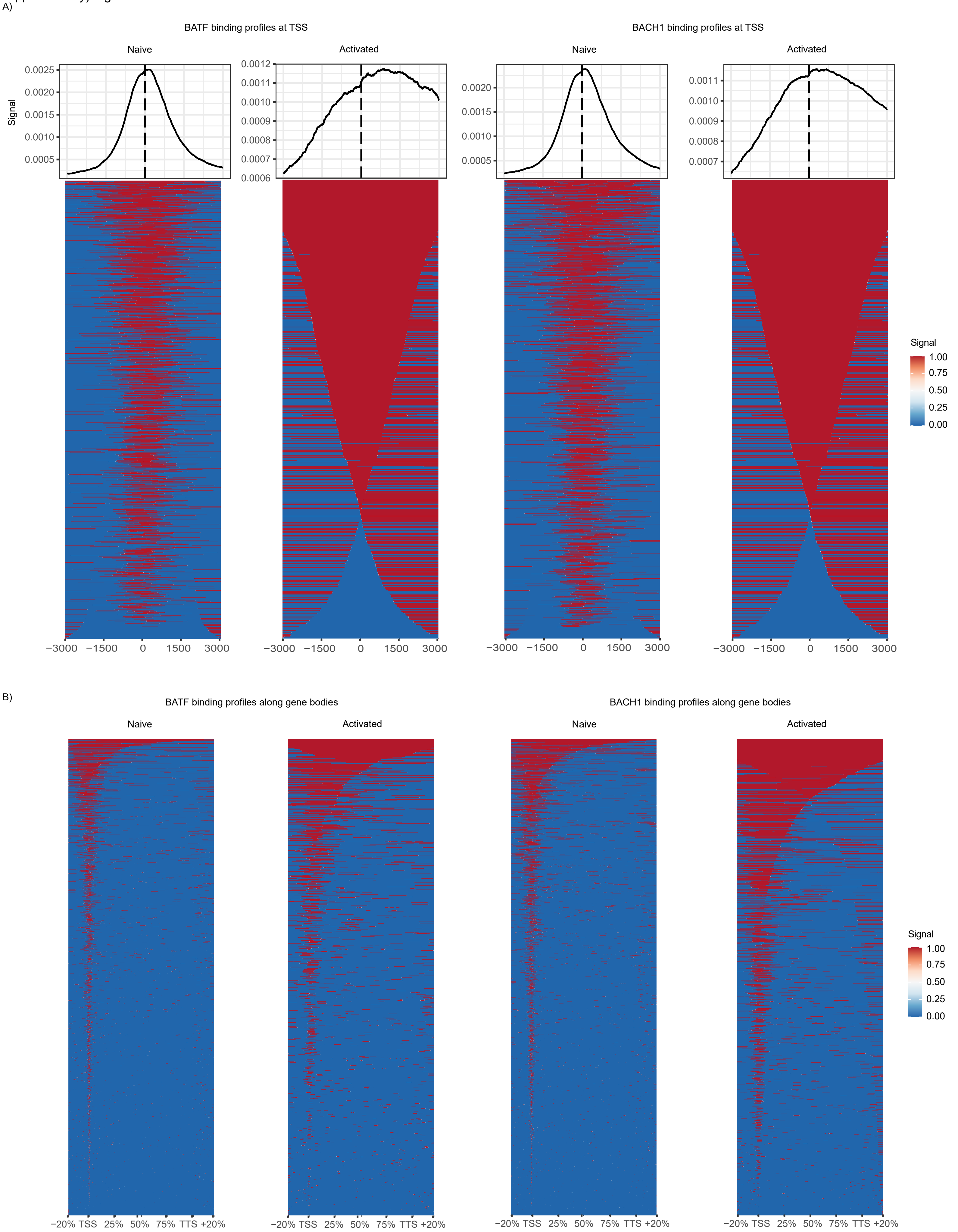

(Supplementary) Figure S14

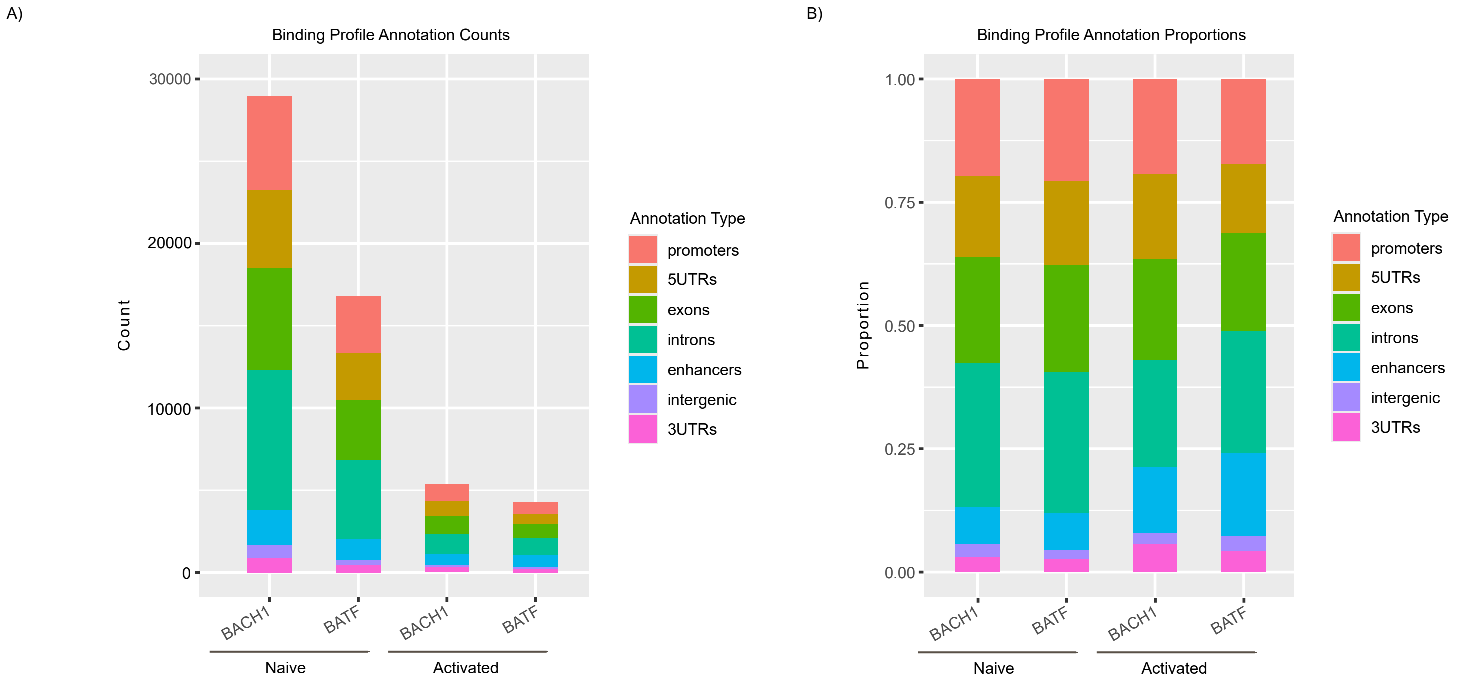
